# Evolution of host-microbe cell adherence by receptor domain shuffling

**DOI:** 10.1101/2021.08.24.457561

**Authors:** EmilyClare P. Baker, Ryan Sayegh, Kristin M. Kohler, Wyatt Borman, Claire K. Goodfellow, Eden R. Brush, Matthew F. Barber

## Abstract

Stable adherence to epithelial surfaces is required for colonization by diverse host-associated microbes. Successful attachment of pathogenic microbes via surface adhesin molecules is also the first step in many devastating infections. Despite the primacy of epithelial adherence in establishing host-microbe associations, the evolutionary processes that shape this crucial interface remain enigmatic. Carcinoembryonic antigen associated cell adhesion molecules (CEACAMs) encompass a multifunctional family of vertebrate cell surface proteins which are recurrent targets of bacterial surface adhesins at epithelial surfaces. Here we show that multiple members of the primate CEACAM family exhibit evidence of repeated natural selection at protein surfaces targeted by bacteria, consistent with pathogen-driven evolution. Inter-species diversity of CEACAM proteins, between even closely-related great apes, determines molecular interactions with a range of bacterial adhesins. Phylogenetic analyses reveal that repeated gene conversion of CEACAM extracellular domains during primate divergence plays a key role in limiting bacterial adhesin tropism. Moreover, we demonstrate that gene conversion has continued to shape CEACAM diversity within human populations, with abundant CEACAM1 variants mediating evasion of adhesins from *Neisseria gonorrhoeae*, the causative agent of gonorrhea. Together this work reveals a mechanism by which gene conversion shapes first contact between microbes and animal hosts.

## INTRODUCTION

Epithelial surfaces are typically the initial point of contact between metazoans and microbes (Brown and Clarke, 2017). As such, host factors at this barrier play an important role in facilitating or deterring microbial colonization. Bacterial attachment to epithelial surfaces is often mediated by a broad class of surface proteins termed adhesins (Kline et al., 2009). In addition to permitting the growth and colonization of commensal microbes, adhesins are also key virulence factors for many pathogenic bacteria. Adhesin-mediated adherence to host cells is often required for other downstream processes including biofilm formation, epithelial invasion, and the delivery of toxic effectors into host cells (Kline et al., 2009; Sadarangani et al., 2011) (Figure 1). Microbial adherence can also trigger epithelial cell signaling cascades, further shaping host responses to resident and invasive microbes. Despite the fundamental importance of epithelial adherence for bacterial colonization and infectious disease pathogenesis, the dynamics of these interactions between host surface proteins and bacterial adhesions over evolutionary timescales remain a mystery. Theory predicts that exploitation of host proteins by pathogens places a significant burden on host populations, driving selection for beneficial mutations in these proteins that limit microbial invasion or virulence.

**Figure 1.**
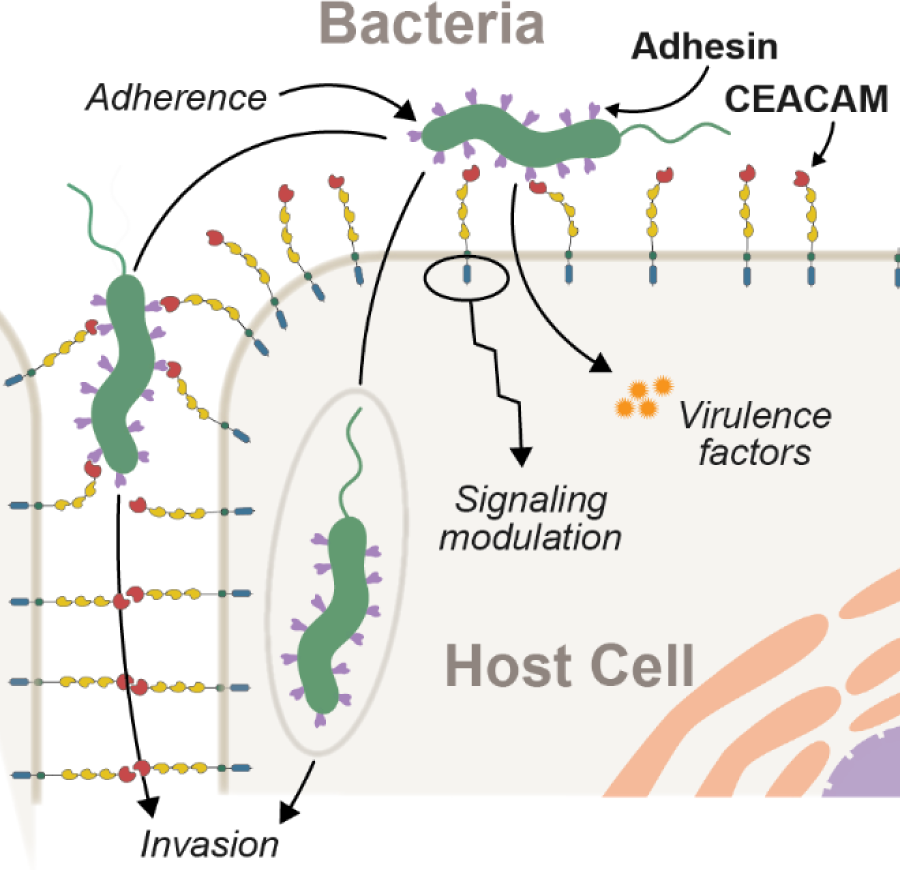
Interactions between epithelial CEACAMs and bacterial adhesins. Bacterial attachment to host cells via adhesin proteins (purple) facilitates epithelial adherence. Adhesins also contribute to pathogenicity by promoting invasion, modulation of host cell signaling pathways, and by promoting the delivery of virulence factors into the host cell cytoplasm.

From a microbial perspective, host defenses can also pose an existential threat resulting in reciprocal adaptation to enhance colonization, growth, and transmission. These cycles of conflict can lead to so-called Red Queen dynamics where each population must continuously adapt simply to maintain its relative fitness (Aleru and Barber, 2020; Brockhurst et al., 2014; Hamilton et al., 1990; Van Valen, 1973). However, pathogens hijack many host factors not directly involved in immunity, possibly limiting their adaptive potential in response to pathogen interaction. For example, epithelial surface proteins are not only essential for interacting with the environment but also serve crucial cellular and physiological functions including barrier maintenance, cell-cell communication, as well as coordinating host physiological and developmental pathways (Kuespert et al., 2006). Consequently, it remains unclear the extent to which such proteins are able to adapt in the face of pathogen antagonism.

A major target of bacterial adhesins on vertebrate epithelia are the carcinoembryonic antigen-related cell adhesion molecule (CEACAM) family of proteins (Gray-Owen and Blumberg, 2006). Collectively, CEACAMs are expressed on nearly all vertebrate epithelial surfaces including the microbe-rich surfaces of the urogenital, respiratory, and gastrointestinal tracts. Epithelial CEACAMs play a variety of roles in cell adhesion as well as intra- and intercellular signaling (Gray-Owen and Blumberg, 2006; Kuespert et al., 2006; Tchoupa et al., 2014). A subset of CEACAMs are also expressed on other cell types, including T-cells and neutrophils where they play important roles in immune signaling and pathogen recognition. CEACAMs typically consist of an extracellular N-terminal IgV-like domain (also termed the N-domain), a variable number of IgC-like domains, and either a membrane anchor or a cytoplasmic signaling domain. Protein-protein interactions involving CEACAMs have been shown to primarily occur through the N-domain (Kuespert et al., 2007; Markel et al., 2004). While the functions of many CEACAM proteins remain obscure, mammalian CEACAM1, CEACAM5 (also known as CEA), and CEACAM6 have been shown to contribute to immunoregulation, cell- cycle progression, and development (Gray-Owen and Blumberg, 2006; Kuespert et al., 2006; Tchoupa et al., 2014).

A growing number of bacterial genera have been found to target CEACAM proteins to promote epithelial adherence and host colonization, including *Neisseria*, *Haemophilus*, *Escherichia*, *Fusobacterium*, *Streptococcus*, and *Helicobacter* (Brewer et al., 2019; Gray-Owen and Blumberg, 2006; Javaheri et al., 2016; Königer et al., 2016; van Sorge et al., 2021). The distinct protein structures and binding mechanisms of these adhesins indicates that CEACAM recognition has arisen independently multiple times during bacterial evolution. While capable of causing serious infections, many of the bacteria that bind CEACAMs also colonize the host as benign commensals. Bacterial CEACAM recognition can lead to several distinct outcomes (Figure1). First, adherence to epithelial CEACAMs can provide a stable habitat to support bacterial growth and proliferation. In mice, for example, expression of human CEACAM1 is sufficient to establish stable colonization by otherwise human-restricted strains of *Neisseria meningitidis* (Johswich et al., 2013). Second, CEACAM binding may facilitate bacterial dissemination through the host epithelium (Wang et al., 1998). Third, in the case of the bacterium *Helicobacter pylori*, CEACAM-adhesin interactions promote the translocation of virulence factors into host cells via the type 4 secretion system (T4SS) leading to severe gastritis and stomach ulcers in humans (Javaheri et al., 2016; Königer et al., 2016). Finally, bacterial adhesins can potentiate CEACAM mediated signaling cascades to manipulate cellular functions, including preventing immune cell activation (Gur et al., 2019a, 2019b; Sadarangani et al., 2011), increasing cellular adhesion to prevent shedding of infected cells (Muenzner et al., 2016, 2010), and activation of apoptosis (Dje N’Guessan et al., 2007).

Previous work has indicated that mammalian CEACAMs have undergone repeated gene gain and loss as well as experienced high levels of sequence divergence (Adrian et al., 2019; Gibbs et al., 2007; Kammerer and Zimmermann, 2010; Pavlopoulou and Scorilas, 2014). These findings, coupled with the observation that many CEACAM-binding bacteria possess a narrow host range, suggests that host genetic variation may be a major determinant of bacterial colonization. In the case of CEACAM3, which is expressed exclusively in neutrophils and aids in destruction of CEACAM-binding bacteria, there is compelling evidence that residues at the interface of adhesin binding are evolving rapidly in a manner consistent with positive selection (Adrian et al., 2019). However, the consequences of epithelial CEACAM evolution for microbial interactions remain unclear. In this study, we investigate patterns of CEACAM divergence in primates and propose how CEACAM evolution and human polymorphisms have shaped interactions with pathogenic bacteria.

## RESULTS

### The CEACAM gene family exhibits repeated episodes of positive selection in primates

To assess patterns of primate CEACAM gene evolution, we compiled sequences of human CEACAM orthologs present in publicly available genome databases. In total nineteen representative species were analyzed including four New World monkeys, ten Old World monkeys, and five hominid species (see Materials and Methods). Some orthologs of human CEACAMs were not identified in a subset of primate genomes, likely due to losses or gains of specific CEACAMs along different lineages or incomplete genome assembly. With the exception of CEACAM3, for which additional exons annotated in Old World monkeys were included (detailed in Materials and Methods), only genomic sequences that aligned to annotated human exons were used for subsequent phylogenetic analyses. To determine if primate CEACAMs have been subject to positive selection, protein-coding sequences were analyzed using the PAML NS sites package (Yang, 2007). This program uses a maximum likelihood framework to estimate the rate of evolution of each gene or codon, expressed as the ratio of normalized nonsynonymous (dN) to synonymous (dS) nucleotide substitutions (dN/dS or ⍵), under different models of evolution. An excess of nonsynonymous substitutions relative to synonymous substitutions between orthologs can suggest that beneficial mutations have been repeatedly fixed by positive selection. A comparison of models that allow and disallow sites evolving under positive selection (⍵ > 1) can determine the likelihood that a particular protein coding sequence has been evolving under positive selection. We found that eight of the twelve primate CEACAM paralogs in our dataset possess genetic signals of positive selection (p-value ≤ 0.05; Supplementary file 1) including CEACAM1, CEACAM3, CEACAM5 and CEACAM6 which have previously been shown to interact with bacterial adhesins (Gray-Owen and Blumberg, 2006). In addition, we also identified elevated ⍵ values for CEACAM7, CEACAM8, CEACAM18 and CEACAM20.

To identify specific amino acid positions that contribute to signatures of positive selection, we analyzed CEACAM sequences using the Bayes Empirical Bayes analysis as implemented in the PAML NS sites package, as well as the programs FUBAR and MEME from the HyPhy software package (Supplementary file 2). To control for the potential impact of recombination on these inferences, we used the program GARD to identify potential breakpoints in our datasets and perform phylogenetic analyses using GARD-informed phylogenies for separate gene segments. Our analyses collectively revealed that sites with elevated ⍵ were concentrated in the N-domain of many CEACAM proteins (Figure 2A; Figure 2 – figure supplement 1). Sites under positive selection in CEACAM18 and CEACAM20 were more dispersed throughout the protein, not localizing to a specific domain. The statistical support for positive selection of CEACAM18 and CEACAM20 in primates was also modest compared to that for other CEACAM proteins.

**Figure 2.**
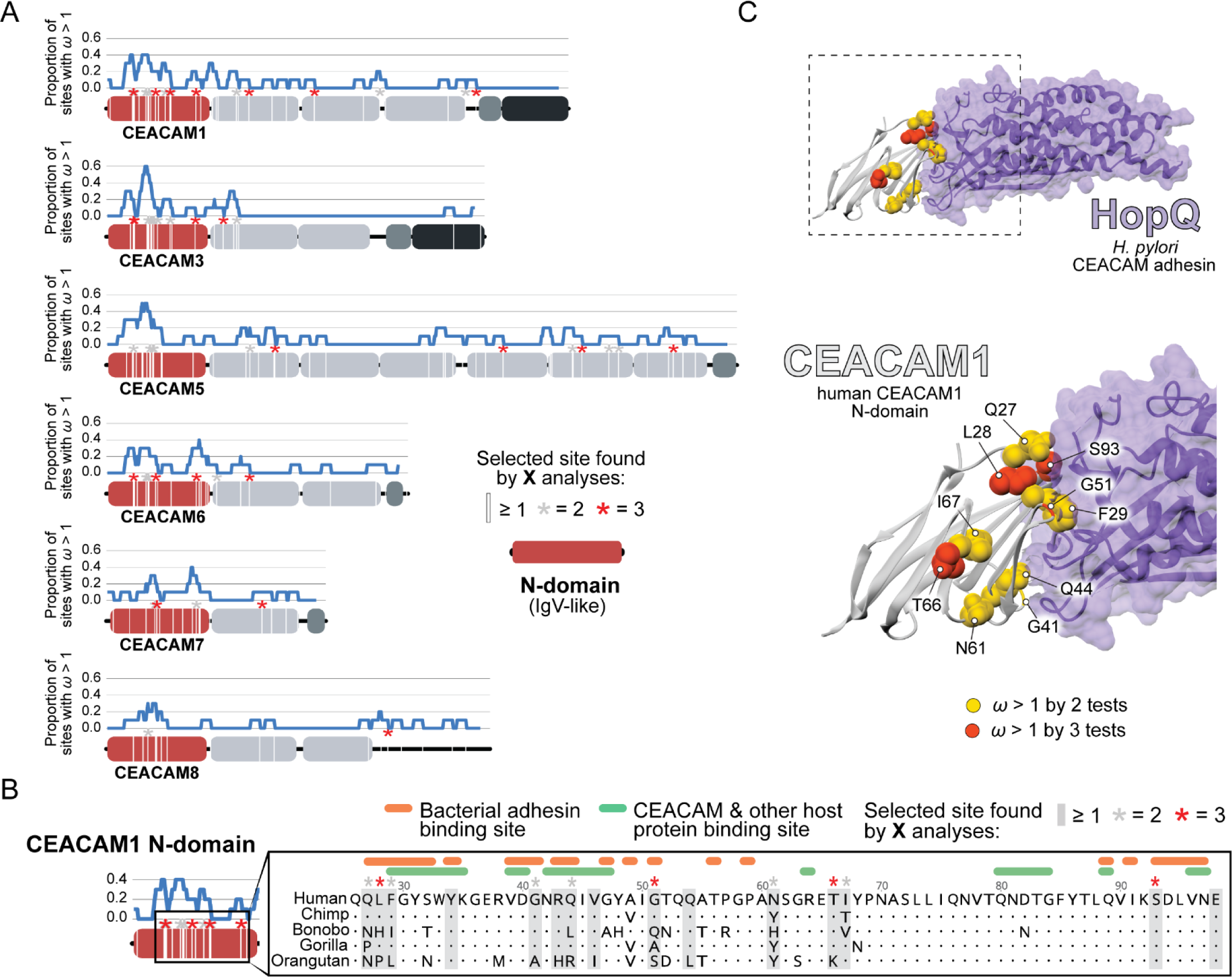
Rapid evolution of primate CEACAM N-domains. A) Sites in CEACAM proteins exhibiting elevated ⍵. Domain structure of CEACAMs outlined in red (N-domain) and gray (all other domains). All rapidly evolving sites identified by at least one phylogenetic analysis (PAML, FUBAR, or MEME) are marked by a white line, sites identified by two or three tests signified by gray and red asterisks respectively. Blue line shows the proportion of rapidly evolving sites identified across a ten amino acid sliding window. B) Multiple sequence alignment of hominid CEACAM1 residues 26-98. Sites identified as evolving under positive selection and sites known to influence adhesin and host protein binding are highlighted (Supplementary file 3). C) Protein co-crystal structure of human CEACAM1 and the HopQ adhesin from H. pylori strain G27 (PDB ID: 6GBG). CEACAM1 sites identified as evolving under positive selection by two or more tests highlighted. **Figure supplement 1**. Primate CEACAM evolutionary analysis summary.

We next sought to determine the functional impact of divergence at rapidly evolving sites in the CEACAM N-domain. Residues that contribute to protein-protein interactions have been extensively annotated for CEACAM1, involving both host factors and bacterial adhesins. Overlaying sites under positive selection with known adhesin and host protein binding sites (Supplementary file 3) revealed extensive overlap between all three categories (Figure 2B) and demonstrates that sites with elevated ⍵ tend to cluster on the protein binding surface. Mapping rapidly-evolving CEACAM1 residues onto a co-crystal structure of human CEACAM1 and the HopQ adhesin from *H. pylori* (Moonens et al., 2018), a known interaction partner, confirmed that multiple sites fall along the binding interface of the two proteins (Figure 2C). In summary, these results demonstrate that multiple primate CEACAM orthologs exhibit signatures of repeated positive selection within the N-domain which facilitates bacterial and host protein interactions.

### CEACAM divergence in primates impairs recognition by multiple bacterial adhesins

To assess how rapid divergence of primate CEACAMs influences recognition by bacterial adhesins, we focused on CEACAM1 which is widely-expressed across different cell types (Gray-Owen and Blumberg, 2006) and has numerous well-documented microbial interactions (Supplementary file 3). Recombinant GFP- tagged CEACAM1 N-domain proteins from a panel of primate species were expressed and purified from mammalian cells (see Materials and Methods). Previous studies have demonstrated that the CEACAM N- domain is both necessary and sufficient to mediate interactions with bacterial adhesins (Javaheri et al., 2016; Kuespert et al., 2007; Markel et al., 2004). We focused our experiments on CEACAM1 binding to two distinct classes of bacterial adhesins: HopQ encoded by *Helicobacter pylori*, and the Opa family adhesins expressed by *Neisseria* species.

The HopQ adhesin is a *H. pylori-*specific outer membrane protein that appears to be universally encoded by *H. pylori* strains and whose interaction with human CEACAM1 has been well-characterized (Bonsor et al., 2018; Javaheri et al., 2016; Königer et al., 2016; Moonens et al., 2018). For our assays we used the common *H. pylori* laboratory strains G27 (Baltrus et al., 2009), J99 (Alm et al., 1999), and Tx30a (ATCC® 51932), which have previously been confirmed to bind human CEACAM1 (Javaheri et al., 2016). The HopQ proteins encoded by these strains encompass the two major divisions of HopQ diversity, termed Type I and Type II (Cao and Cover, 2002; Javaheri et al., 2016). Strains G27 and J99 both encode a single copy of a Type I HopQ adhesin, while Tx30a encodes a Type II HopQ adhesin. All strains include extensive divergence in the CEACAM1 binding region (Bonsor et al., 2018; Moonens et al., 2018). Opa proteins are a highly diverse class of adhesins encoded by *Neisseria* species that are structurally distinct from the HopQ adhesin (Bonsor et al., 2018; Fox et al., 2014; Moonens et al., 2018; Sadarangani et al., 2011). Despite their limited sequence identity, both Opa52 and Opa74 are known to bind human CEACAM1 (Roth et al., 2013). Because *Neisseria* species typically encode multiple unique phase-variable Opa variants, individual Opa genes from *N. gonorrhoeae* were cloned and expressed heterologously in K12 *Escherichia coli*, which does not bind to CEACAM proteins.

To assess pairwise interactions between primate CEACAMs and bacterial adhesins, we incubated recombinant CEACAM1 N-domain proteins with individual bacterial strains. Bacterial cells were washed, pelleted, and the presence of bound CEACAM1 protein was assessed by western blot. We observe that all bacterial strains tested bind to the human CEACAM1 N-domain, consistent with previous studies (Figure 3A). Incubation of *H. pylori* strain G27 with GFP alone fails to yield detectable signal, confirming that binding is CEACAM-dependent (Fig 3B). Furthermore, a *Δhopq* mutant of strain G27 does not exhibit significant CEACAM1 binding, consistent with previous reports that HopQ is the sole CEACAM-binding adhesin present in these strains (Figure 3 – figure supplement 1).

**Figure 3.**
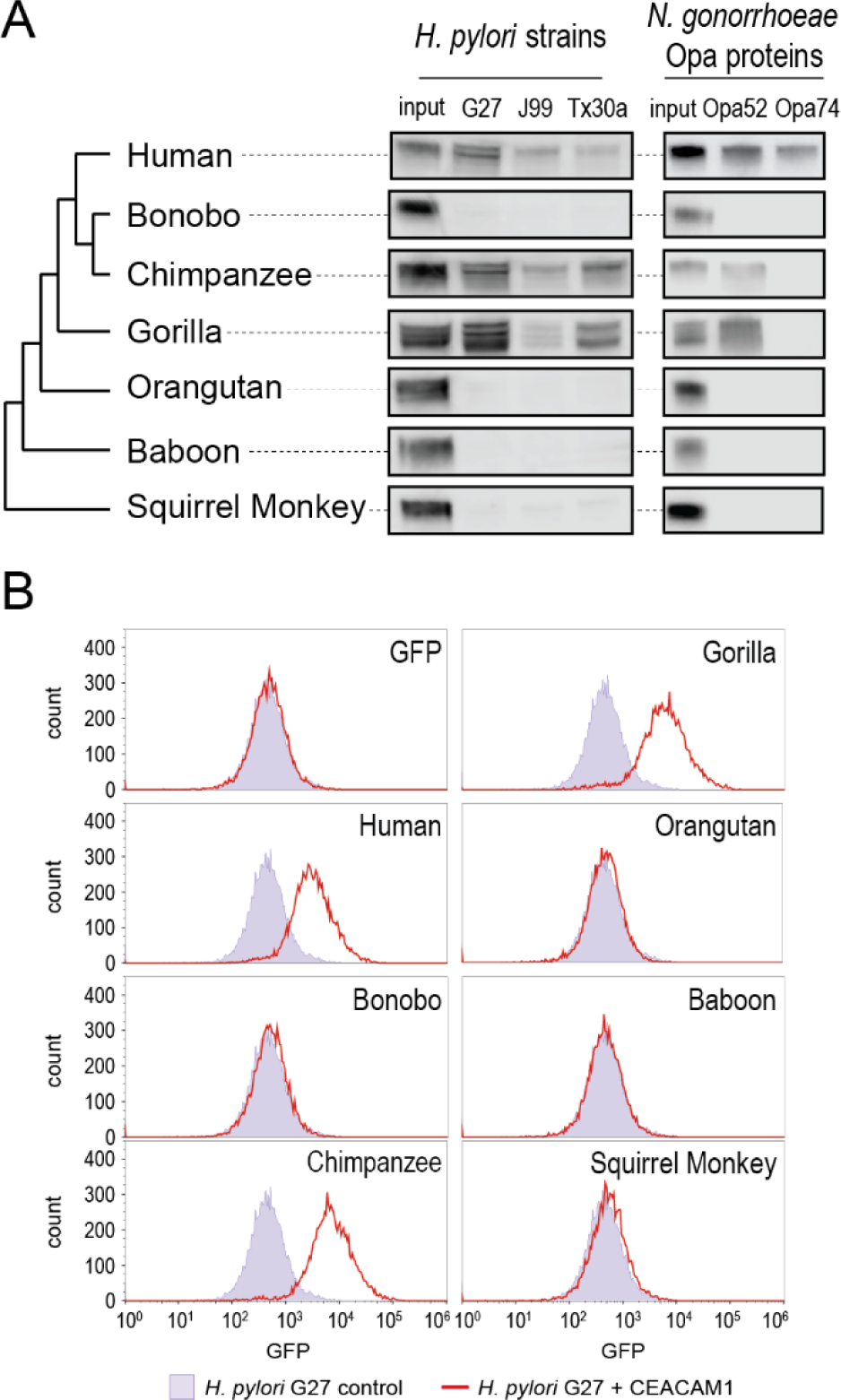
CEACAM1 divergence in great apes restricts bacterial adhesin recognition. A) Binding between primate GFP tagged CEACAM1 N-domain orthologs and bacteria determined by pulldown assays and visualized by western blotting. Input is 10% CEACAM1 protein used in bacterial pulldowns. Primate species relationships indicated by phylogenetic tree. B) Pulldown experiments of *H. pylori* strain G27 incubated with CEACAM1 N-domain constructs or GFP alone assayed by flow cytometry. Binding indicated by GFP fluorescence. Representative western blot and flow cytometry experiments depicted. **Source data 1.** Raw and labeled western blot images for Figure 3A and flow cytometry data for Figure 3B. Figure supplement 1. *H. pylori* G27 *Δhopq* pulldown Figure supplement1 – source data 1. Raw and labeled western blot images for Figure 3 – figure supplement 1

Examining non-human CEACAM1 bacterial binding, the chimpanzee CEACAM1 N-domain, which differs from the human protein at four amino acid positions, binds to all adhesin-expressing strains except Opa74. Gorilla CEACAM1, which differs from the human N-domain at five sites (three non-overlapping with chimpanzee) is also unable to bind Opa74 but does bind *H. pylori* strains and Opa52. Orangutan CEACAM1 is unable to interact with any bacterial strains, nor do baboon and squirrel monkey. We noted that despite the limited species divergence between bonobos and chimpanzees, bonobo CEACAM1 does not bind any of the tested bacterial strains (Figure 3A). Previous studies have found the results of CEACAM-binding assays to be consistent between western blotting and by flow cytometry (Adrian et al., 2019; Javaheri et al., 2016; Königer et al., 2016; Kuespert et al., 2007). We confirmed this for our system with *H. pylori* strain G27, using flow cytometry to detect specific binding of GFP-tagged CEACAMs on the bacterial cell surface (Figure 3B). These results demonstrate that CEACAM1 N-domain divergence between closely-related primate species, even within the great apes, determines bacterial recognition in an adhesin-specific manner.

### Recurrent gene conversion of primate CEACAM N-domains

The inability of *H. pylori* strains or *N. gonorrhoeae* adhesins to bind bonobo CEACAM1 was surprising given bonobo’s close phylogenetic relationship to both humans and chimpanzees. While archaic humans are believed to have diverged from our primate relatives at least 5 million years ago, the major divergence between chimpanzees and bonobos occurred only one to two million years ago (Prado-Martinez et al., 2013). Closer inspection revealed that the bonobo CEACAM1 N-domain sequence is unusually divergent from that of both humans and chimpanzees, while other regions of the coding sequence show higher degrees of identity (Figure 4 – figure supplement 1A). To investigate bonobo CEACAM1 evolution further, we first validated the bonobo CEACAM1 N-domain sequence present in our bonobo reference genome through comparison of assemblies and sequencing reads from multiple bonobo individuals (see Materials and Methods). Having confirmed the identity of the bonobo CEACAM1 reference sequence, we compared this gene to sequences from other hominids. Relative to its orthologs in humans and chimpanzees, bonobo CEACAM1 differs at nearly 20% of sites in the N-domain whereas humans and chimpanzees differ at only about 4% of sites. In contrast, outside of the N-domain bonobo CEACAM1 diverges from humans and chimpanzees at approximately 2% of sites, while human and chimpanzee CEACAM1 differ at around 1% of sites. We also noted that the number of divergent sites between bonobo and human in the N-domain (18 residues) is nearly identical to the number of divergent sites between bonobo and chimpanzee (20 residues), despite the closer phylogenetic relationship between bonobos and chimpanzees. In fact, the divergence between the bonobo and chimpanzee CEACAM1 N-domains is greater than that between chimpanzee and the earliest diverging member of the hominid clade, orangutan (81% versus 83% amino acid identity respectively). A comparison of N-domain sequences for CEACAM5, another rapidly evolving CEACAM, further highlights the extreme divergence of bonobo CEACAM1. Between human CEACAM5 and the bonobo and chimpanzee CEACAM5 sequences there are only ten and nine amino acid changes respectively, while bonobo and chimpanzee differ at only five sites along the entire length of the N-domain (Figure 4 – figure supplement 1B).

**Figure 4.**
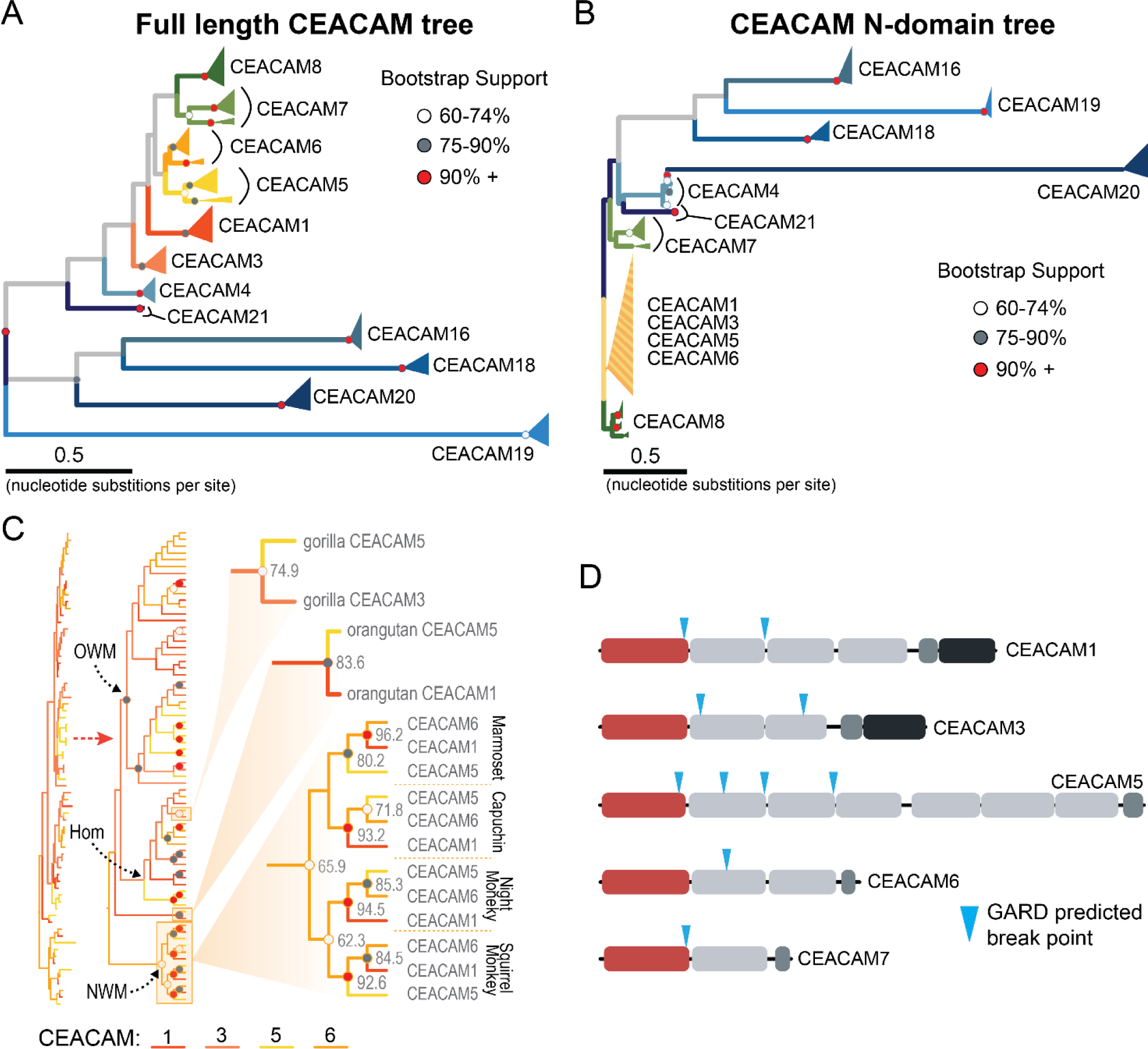
Recurrent episodes of gene conversion among adhesin-binding CEACAMs. A) Maximum-likelihood based phylogenetic reconstruction of full-length primate CEACAM protein coding sequences. B) Phylogenetic reconstruction of the IgV-like (N-domain) of primate CEACAM proteins. C) Expanded view of the clade containing the N-domains of CEACAM1, CEACAM3, CEACAM5 and CEACAM6 from panel B. Arrows indicate nodes designating clades for Old World monkeys (OWM), hominids (Hom) and New World monkeys (NWM). Specific subclades, gorilla CEACAM3 and CEACAM5, orangutan CEACAM5 and CEACAM1, and New World monkeys are further magnified and highlighted with bootstrap support at nodes. D) Domain structures of CEACAM proteins predicted to have undergone recombination by GARD analysis with sites of predicted breakpoints highlighted (blue arrows). CEACAM N-domains are denoted in red. Figure 4 *legend continued on next page*. Figure supplement 1. Alignment of Human-Pan CEACAM sequences. Figure supplement 2. Expanded full length CEACAM tree Figure supplement 3. Expanded CEACAM N-domain tree. Figure supplement 4. Expanded view of CEACAM1,3,5,6 N-domain clade. Figure supplement 5. Expanded CEACAM IgC domains tree. Figure supplement 6. Expanded CEACAM cytoplasmic domain tree.

The degree of divergence within the N-domain of bonobo CEACAM1 suggests processes other than sequential accumulation of single nucleotide mutations could be responsible. One mechanism by which this could occur is through gene conversion, a form of homologous recombination in which genetic material from one location replaces sequence in a non-homologous location, often with substantial sequence similarity (Chen et al., 2007). Gene conversion is thought to be an important source of genetic novelty and a mechanism that can accelerate adaptation (Bittihn and Tsimring, 2017; Daugherty and Zanders, 2019). To determine if inter-locus recombination has shaped the evolution of CEACAM genes in primates, we looked for evidence of discordance between species and gene trees. Gene-species tree discordance can be an indication of multiple evolutionary processes, including a history of gene conversion between paralogs. In a maximum likelihood-based phylogeny of full-length CEACAM coding sequences, clades containing single CEACAM paralogs were inferred with robust statistical support (Figure 4A, Figure 4 – figure supplement 2). In general, the relationships between CEACAM homologs are inferred with high confidence and reflected species relationships as expected for the divergence of orthologous coding sequences. To determine if there have been domain-specific instances of gene conversion, we constructed phylogenetic trees of specific

CEACAM domains. Typically, we expect paralog sequences to form clearly defined clades reflecting species divergence. This is the pattern we observe for full-length CEACAM coding sequences, indicating that overall the paralogs have remained distinct since their initial duplication and have steadily diverged between species. Specific CEACAM domain sequences generally follow this pattern (Figure 4B, Figure 4 – figure supplement 3-6). However, the N-domains of CEACAM1, CEACAM3, CEACAM5 and CEACAM6 deviate strikingly from this norm and form a single monophyletic group (hereafter called CCM1356), albeit one with low bootstrap support (Figure 4B, Figure 4 – figure supplement 3 and 4). Within the CCM1356 clade we observe that rather than clustering by paralog, N-domains are split into subclades representing the three major primate lineages (Figure 4C, Figure 4 – figure supplement 4). In general, the close phylogenetic relationship of sequences within these clades is well-supported. This topology suggests that these CEACAM N-domains are more similar to paralogous domains within the same species or primate lineage than they are to their respective orthologs across species. Several well-supported nodes provide further evidence that gene conversion is driving concerted evolution within the CCM1356 clade (Figure 4C). Certain pairs of N-domains, such as CEACAM3 and CEACAM1 in gorilla and CEACAM1 and CEACAM5 in orangutan, form monophyletic groups with strong bootstrap support. As these relationships are not observed for the other domains of these CEACAM proteins, this suggests conversion events affecting only the N-domains of these CEACAMs occurred in these species. New World monkeys provide the most striking phylogenetic evidence of gene conversion among primates. For each of the four New World monkey species examined, the N-domains of CEACAM1, CEACAM5, and CEACAM6 are all more closely related within species than to their orthologs in other species, suggesting gene conversion has independently acted on the N-domains of these three CEACAMs at least four times within this single clade (Figure 4C). These findings are consistent with N- domains of CEACAMs 1, 3, 5, and 6 undergoing widespread concerted evolution, likely facilitated by gene conversion.

To further test for evidence of gene conversion acting on primate CEACAM family members, we applied the GARD algorithm from the HyPhy software package. GARD detects topological changes between trees inferred from segments of a gene alignment, assesses the likelihood they are consistent with recombination, and identifies potential breakpoints. Consistent with our phylogenetic examination of CEACAM homologs, GARD detects strong evidence of recombination for CEACAM1, CEACAM3, CEACAM5 and CEACAM6 (Figure 4D). In all cases, breakpoints were identified at the C-terminus of the N-domain or in immediately adjacent IgC domains. This pattern is consistent with repeated N-domain gene conversion between CCM1356 paralogs (Figure 4D) and is also in line with our phylogenetic reconstructions of CEACAM IgC domains (Figure 4 – figure supplement 5). In addition to CEACAM1, CEACAM3, CEACAM5, and CEACAM6, GARD also indicates a recombination breakpoint for CEACAM7 that would encompass the N-domain. While we do not detect discordance in our N-domain gene tree that implicates gene conversion involving CEACAM7, there is a single instance in the IgC domain tree of a gorilla CEACAM5 IgC domain grouping more closely with homologs of the IgC domain of CEACAM7 (Figure 4 – figure supplement 5). A breakpoint in this region is also consistent with CEACAMs with rapid N-domain evolution being involved in gene conversion events as well as previous analyses (Zid and Drouin, 2013). Together these results support a model in which gene conversion between rapidly diverging CEACAMs has contributed to N-domain diversification during primate evolution.

### Rapidly evolving regions of CEACAM1 are sufficient to block bacterial adhesin recognition

Phylogenetic analyses confirm that the bonobo CEACAM1 N-domain is not closely related to other primate CEACAM1 sequences but fail to strongly support its relationship to any other single CEACAM N- domain. Reasoning that the extant bonobo CEACAM1 gene may have arisen from multiple iterative recombination events, we performed a BLAST search of genomes on the NCBI database using base pairs 103-303 of the bonobo CEACAM1 sequence (corresponding to resides 1-67 of the N-domain) as our query. Human and chimpanzee are roughly 86% identical to bonobo CEACAM1 in this region versus 99% identical (a single nucleotide change) in the remaining 120 base pairs (Figure 4 – figure supplement 1). This search identifies orangutan CEACAM3 as the closest match. While the similarity between the first 120bp of bonobo CEACAM1 and orangutan CEACAM3 is striking and the final third of the nucleotide sequence is nearly identical to human and chimpanzee CEACAM1, other segments of bonobo CEACAM1 are still quite divergent from all other N-domain sequences (Figure 5A). A BLAST search of this region in bonobo CEACAM1 (base pairs 221-380) indicates the greatest similarity is with the analogous region from rhesus macaque CEACAM6.

**Figure 5.**
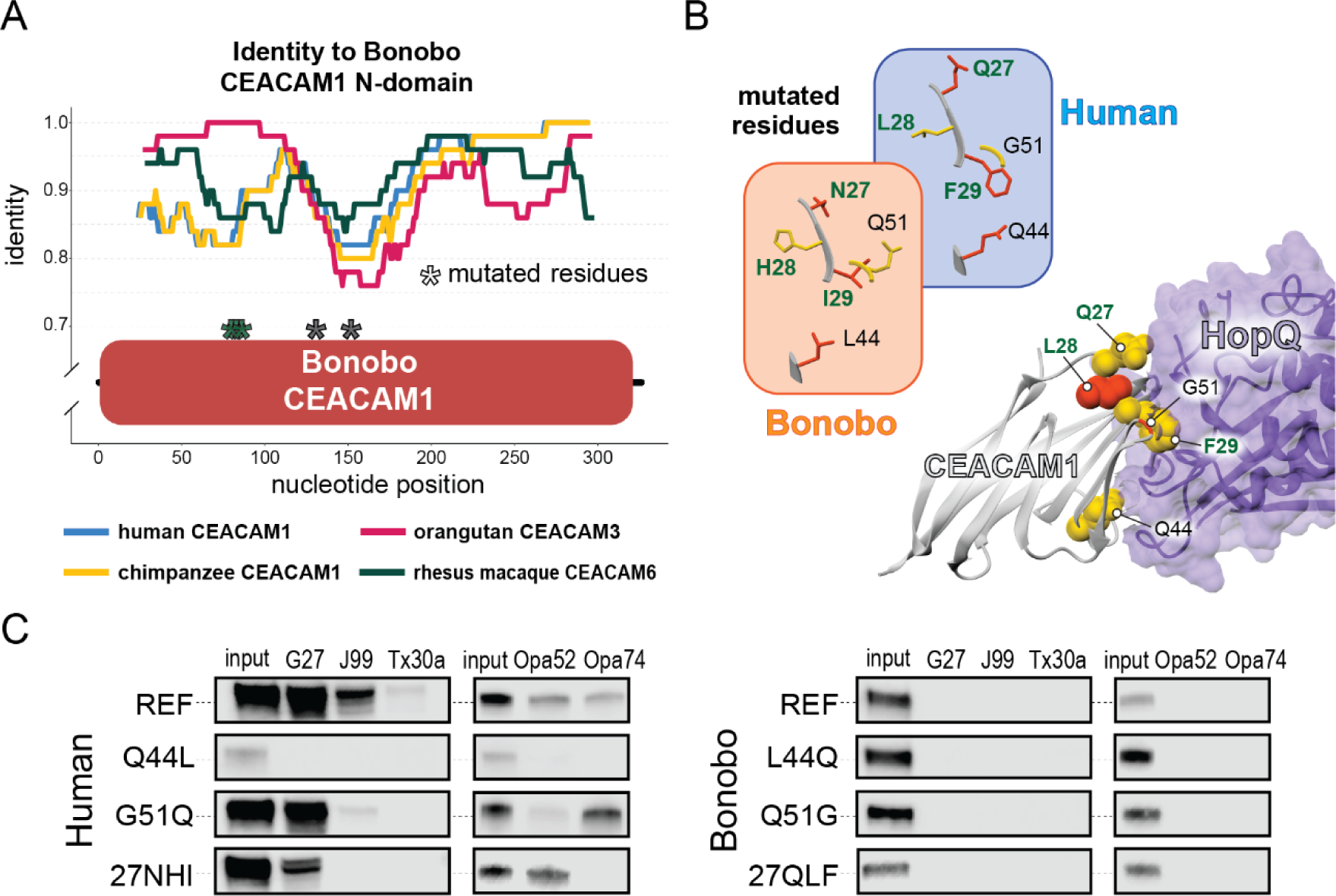
Rapid divergence of the bonobo CEACAM1 N-domain impairs bacterial adhesin recognition. A) Graph shows a fifty base pair sliding window of identity between bonobo CEACAM1 N-domain sequence and other CEACAM sequences. Asterisks mark locations of residues mutated for adhesin binding assays. B) Windows show amino acids and their structures at sites selected for mutational analysis in humans and bonobos. Lower right is a protein co-crystal structure of human CEACAM1 and *H. pylori* G27 HopQ with sites selected for mutagenesis highlighted. C) Representative western blots of pulldown experiments assaying binding between chimeric human and bonobo CEACAM1 N-domain constructs and bacterial strains. **Source data 1.** Raw and labeled western blot images for Figure 5C. Figure supplement 1. Alignment of rapidly evolving N-domain region in hominids.

However, the increased similarity of macaque CEACAM6 in this region compared to other CEACAMs is marginal (Figure 5A).

The extreme divergence of the bonobo CEACAM1 N-domain from other CEACAM1 homologs in even its closest relatives could indicate that this particular sequence has been evolving independently of other N- domain alleles for a long period of time as a result of balancing selection. This has been observed for other genes involved in host-pathogen conflicts, most notably major histocompatibility complex (MHC) alleles. In this case, we might expect to identify alleles similar to bonobo CEACAM1 currently circulating in other hominid populations, and likewise alleles similar to CEACAM1 sequences observed in humans and chimpanzees may be found in the larger bonobo population. In a search of human genetic variation data available through the International Genome Sample Resource (IGSR) accessed through the Ensembl webserver (www.ensembl.org) there is no evidence for any alleles with similarity to bonobo CEACAM1 circulating within human populations. Searching population data from the Great Apes Genome Project (Prado-Martinez et al., 2013), alleles similar to bonobo CEACAM1 are not found for chimpanzees, gorillas, or orangutans. Likewise, CEACAM1 alleles similar to those found in humans and chimpanzees are not observed in any of the bonobo genomes from the same dataset. Given the information at hand, it is difficult to precisely determine the series of mutational events that produced the bonobo CEACAM1 allele or determine the likely origin point of this allele in the diversification of hominids. However, these results are consistent with multiple independent instances of gene conversion giving rise to bonobo CEACAM1, with subsequent fixation of this haplotype in bonobo populations since their divergence from chimpanzees over the last million years.

Given the large number of residue changes between human and bonobo CEACAM1, we next sought to determine if a subset of rapidly-evolving sites are sufficient to either impair or restore recognition by bacterial adhesins. To test this, we generated CEACAM1 N-domain proteins in which a subset of residues between humans and bonobos were swapped. We focused on sites that are identical in humans and chimpanzees but differ in bonobos and which exhibit high ⍵ across primates, resulting in a total of five tested sites (Figure 5A & B). Of these residues we chose to mutate adjacent amino acids 27-29 as a single group. This patch of sites is highly variable among the rapidly-evolving CEACAMs, particularly CEACAM1, CEACAM3 and CEACAM5 (Figure 5 – figure supplement 1). None of the “humanized” mutants in the bonobo CEACAM1 background were sufficient to confer binding (Figure 5C). In contrast, introduction of bonobo residue 44 into human CEACAM1 (mutation Q44L) prevents binding by *H. pylori* and Opa expressing strains, while introduction of bonobo variable sites 27-29 abolishes binding to Opa74 (Figure 5C). Mutation G51Q has no appreciable impact on binding by *H. pylori* strain G27 or Opa74, but blocks binding by strain Tx30a and reduces binding to J99 and Opa52. Collectively these results reveal that multiple single positions in human CEACAM1 exhibiting signatures of positive selection are sufficient to impair recognition by multiple bacterial adhesins. Moreover, these findings also demonstrate how instances of gene conversion between CEACAM paralogs could serve as large-effect adaptive mutations during conflicts with pathogens.

### Abundant human CEACAM1 polymorphisms impair bacterial recognition

Pervasive evidence of positive selection acting on CEACAMs in primates raises the question as to whether CEACAM variants that evade pathogen recognition are currently segregating in human populations. To explore the existence of human CEACAM variants and their consequences for bacterial interactions, we queried human single nucleotide polymorphism (SNP) and haplotype data for rapidly evolving CEACAM paralogs available from the International Genome Sample Resource accessed through the Ensembl genome browser (see Materials and Methods). We found that variation in the N-domains of CEACAM6, CEACAM7 and CEACAM8 predominantly consists of polymorphisms not shared with other CEACAM proteins and found on isolated haplotypes. In contrast, CEACAM1, CEACAM3 and CEACAM5 N-domain variation is composed primarily of extended haplotypes (Figure 6 – figure supplement 1-4). Furthermore, these extended haplotypes increase similarity between CEACAM1, CEACAM3 and CEACAM5, consistent with possible gene conversion events. Indeed, some haplotypes not only have changes at nonsynonymous sites that increase similarity with these CEACAMs, but also include multiple shared synonymous changes. These observations suggest that gene conversion among CEACAMs has occurred relatively recently and may be ongoing in human populations.

**Figure 6.**
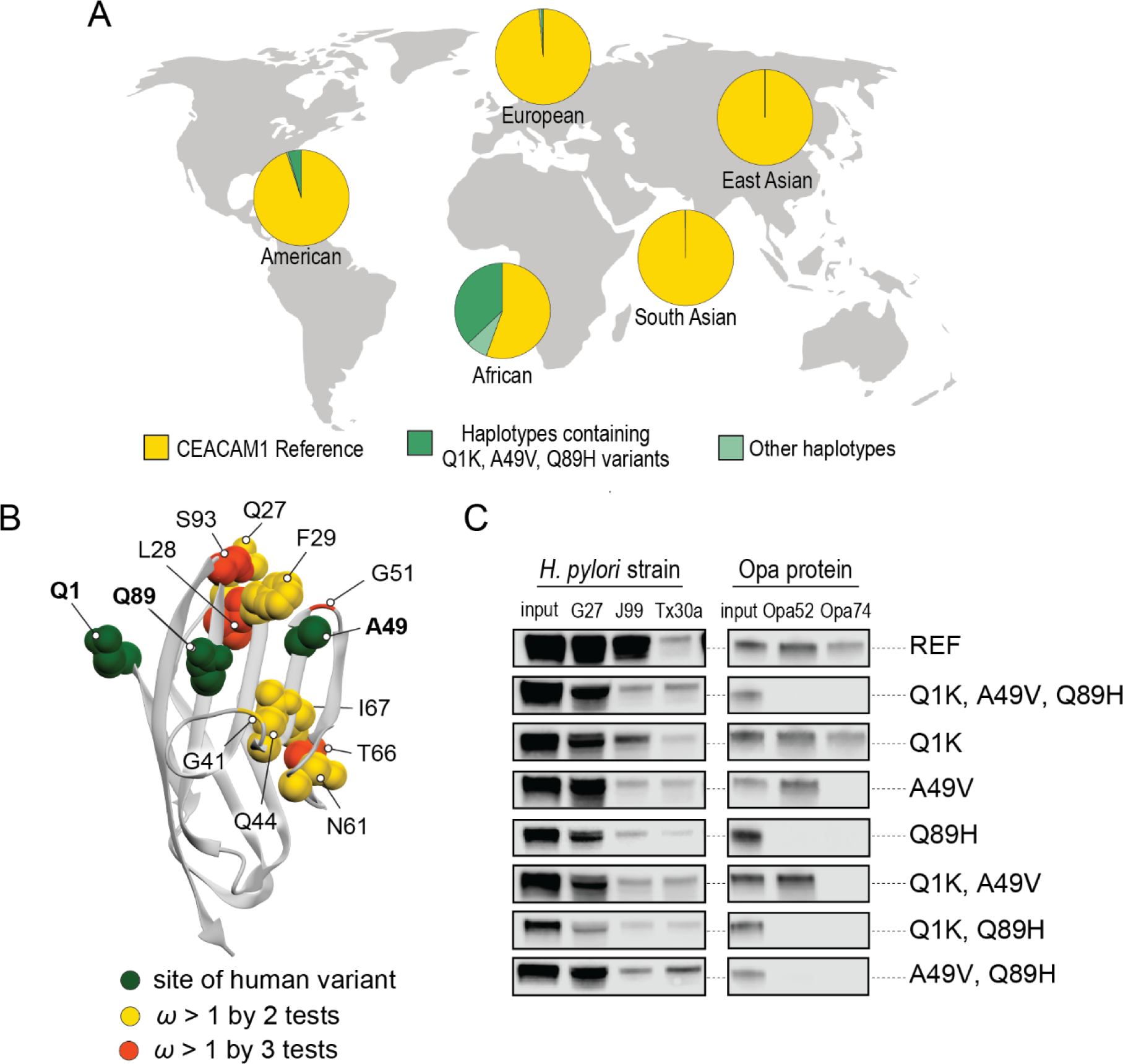
Abundant human CEACAM1 variants restrict pathogen binding. A) Frequency of haplotypes containing high frequency human variants Q1K, A49V and Q89H across human populations (map from <u>BioRender.com</u>). B) CEACAM1 crystal structure highlighting high frequency human variants and sites found to be evolving under positive selection across simian primates. C) Representative western blots of pulldown experiments testing binding between combinations of high frequency human variants in the human CEACAM1 reference background and bacterial strains. D) Sequence identity between the human CEACAM1-CEACAM3 hybrid allele and the human CEACAM1 and CEACAM3 reference alleles. **Source data 1.** Data files and code for analyzing CEACAM1 haplotypes for Figure 6A and Figure 6 –figure supplement 2. Source data 2. Raw and labeled western blot images for Figure 6C. Figure supplement 1. Human CEACAM-like CEACAM1 haplotypes. Figure supplement 2. Human CEACAM1 haplotype frequencies. Figure supplement 3. Human CEACAM3 variation. Figure supplement 3 – source data 1. Data files and code for analyzing CEACAM3 haplotypes for Figure 6 – figure supplement 3B and C. Figure supplement 4. Human CEACAM5 variation. Figure supplement 4 – source data 1. Data files and code for analyzing CEACAM5 haplotypes for Figure 6 – figure supplement 4B and C.

A search of polymorphisms for CEACAM1 in human populations reveals three high-frequency nonsynonymous variants within the N-domain: Q1K (rs8111171), A49V (rs8110904) and Q89H (rs8111468) (Figure 6A). The haplotype containing all three alternative alleles is the most frequent non-reference CEACAM1 haplotype annotated, occurring in 14% of the human population overall and in up to 43% of individuals in African populations (Figure 6A). In total, nearly 17% of all sequenced individuals carry at least one of these high frequency SNPs (Figure 6 – figure supplement 2). Of the three variants, A49V and Q89H both lie within regions of CEACAM1 known to interact with bacterial adhesins suggesting they may alter bacterial adherence (Figure 6B). To determine if these high-frequency CEACAM1 polymorphisms affect bacterial recognition, we generated recombinant CEACAM1 N-domain variant proteins for use in our adhesin binding assays. None of the variants are able to abolish CEACAM1 binding to our panel of *H. pylori* strains (Figure 6C). In contrast, *Neisseria* Opa expressing strains exhibit highly variable recognition of multiple human CEACAM1 variants. The Q1K mutation alone has no impact on binding, while A49V abolishes recognition by Opa74, and variant Q89H abrogates binding to both Opa52 and Opa74 (Figure 6C). Combinatorial CEACAM1 variants reveal that these mutations behave in a dominant manner, with Q89H dominant over A49V (Figure 6C). Together these results demonstrate that high frequency human polymorphisms in CEACAM1 are sufficient to impair binding by specific classes of bacterial adhesins present in human pathogens. These findings further suggest that high-frequency CEACAM variants could alter human colonization or infection by pathogenic *Neisseria*, including causative agents of gonorrhea and meningitis.

## DISCUSSION

Our investigation of species-specific bacterial adherence to CEACAM1 revealed an unforeseen example of extreme genetic divergence within the great apes. The bonobo CEACAM1 gene could represent a rapid succession of single residue changes combined with multiple recombination events arising in bonobos under strong selection and/or a population bottleneck. Alternatively, this allele may be ancient and have been subject to balancing selection or incomplete lineage sorting in ancestral hominid populations. We also considered that the source of the bonobo CEACAM1 sequence may not be from functional CEACAM genes, but a pseudogenized CEACAM sequence or a pregnancy-specific glycoprotein (PSG), a family of proteins closely related to CEACAMs. However, a BLAST search of relevant NCBI databases (see Materials and Methods) fails to identify any new genomic regions in bonobos or other primates with greater sequence identity than what had already been found. While there are multiple possible explanations for the highly divergent nature of bonobo CEACAM1, absent further evidence the origin of this particular allele remains obscure. What is clear from the example of bonobo CEACAM1 however, is the extent to which gene conversion can rapidly generate diversity between closely related species and the impact of such variation on interactions with microbes.

During the course of investigating the origin of the bonobo CEACAM1 sequence we discovered evidence that gene conversion has shaped the evolution of many CEACAMs across primates. While we identify several instances of likely gene conversion, results from phylogenetic analyses probably represent an underestimate of the true number of recombination events that have occurred among rapidly evolving CEACAMs in primates. Repeated episodes of gene conversion can obscure past instances of recombination and hinder their identification by gene-species tree discordance. In turn, GARD analyses and recombination detection programs in general tend to miss many recombination events (Bay and Bielawski, 2011; Kosakovsky Pond et al., 2006). One particularly interesting example in orangutans implicates multiple conversion events impacting CEACAM1, CEACAM5 and CEACAM8. (Figure 5 – figure supplement 1). Phylogenetic analyses indicate a species-specific conversion event between CEACAM1 and CEACAM5 in orangutans. Prior to the CEACAM1-CEACAM5 conversion, however, residues 29-64 in either CEACAM1 or CEACAM5 were likely replaced by the homologous sequence from CEACAM8. Evidence for this event includes not only the multiple nonsynonymous substitutions shared with orangutan CEACAM8, but a shared synonymous substitution in both orangutan CEACAM1 and CEACAM5 only observed in hominid CEACAM8 homologs. Despite this evidence, neither our phylogenetic analyses nor GARD analyses suggest CEACAM8 has been involved in gene conversion. Like for CEACAM7, the involvement of CEACAM8 in intra-paralog gene conversion is consistent with CEACAMs with rapid N-domain evolution participating in gene conversion events. Overall, the rapid shuffling of genetic variation among CEACAM genes that we observe could greatly augment the potential for host adaptation in the face of microbial antagonism.

It has been suggested that gene conversion between CEACAM paralogs works to preserve the ability of CEACAM3 to effectively mimic bacterially antagonized CEACAMs and thereby maintain its function as a decoy receptor (Zid and Drouin, 2013; Zimmermann, 2019). Indeed, our results and those of Adrian *et al*. (Adrian et al., 2019) support the importance of gene conversion for maintaining the similarity of CEACAM3 to other bacterial adhesin binding CEACAMs in apes and Old World monkeys. However, gene conversion cannot only serve to maintain CEACAM3’s mimicry function. There is no evidence that New World monkeys encode a CEACAM3 homolog, yet within this group gene conversion appears to be frequent between CEACAM1, CEACAM5, and CEACAM6 (Figure 4C). Additionally, we observe multiple conversion events in hominids that do not involve CEACAM3 (Figure 4C, Figure 5 – figure supplement 1, and Figure 6 – figure supplement 4).

We propose that gene conversion among epithelial CEACAMs reflects a general mechanism of pathogen evasion (Figure 7), allowing beneficial mutations to be exchanged among CEACAMs that interact with pathogens. Gene conversion allows beneficial sets of mutations to spread, whether between decoys and targets or among antagonized epithelial CEACAMs, more rapidly than conversion of residues through independent mutational events (Bittihn and Tsimring, 2017). In addition, in the case of decoys and targets the ability of sequences to be exchanged back-and-forth limits the residues available for differentiation by pathogens and provides decoys the ability to gain binding to CEACAM antagonists through exchanges from epithelial CEACAMs. Finally, the interchangeability of the protein binding domain among antagonized CEACAMs effectively provides multiple copies of the same domain, increasing the evolutionary space available to CEACAMs to evolve and test alleles, further enhancing the pace at which beneficial alleles may evolve. In this way gene conversion could provide an important mechanism by which the host can keep pace with rapidly evolving pathogenic microbes. It is also notable that the immense variation between Opa alleles, like CEACAMs, has been shown to involve a combination of rapid substitutions and recombination between extracellular loop domains that recognize host factors and serve as potential antigens for the host adaptive immune system. In this regard CEACAM-Opa interactions reflect an unusual evolutionary dynamic in which recombination likely plays a crucial role in reciprocal adaptation.

**Figure 7.**
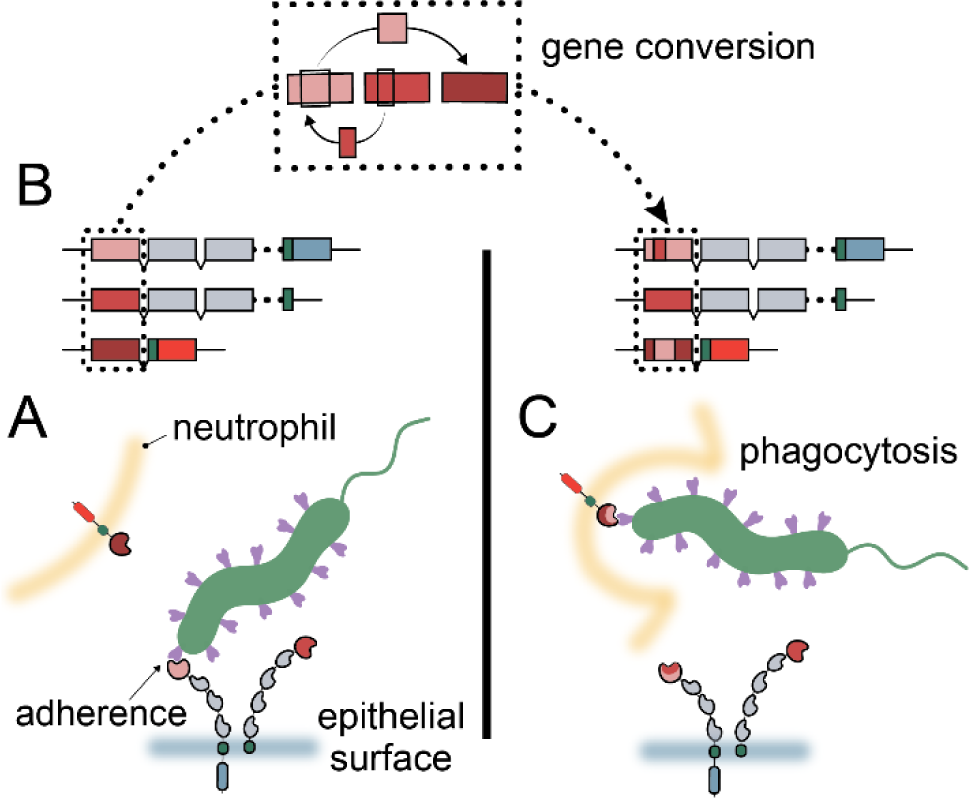
Model of CEACAM evolution in primates. A) Bacterial adhesins recognize a subset of epithelial CEACAM proteins and avoid binding with decoy CEACAM receptors present on neutrophils. B) Gene conversion facilitates the shuffling of regions of the CEACAM N-domain that alter binding to bacterial adhesins. C) Through gene conversion outlined in B, epithelial CEACAM proteins avoid binding by bacterial adhesins while the CEACAM decoy receptor gains binding triggering bacterial clearance through phagocytosis.

In addition to exploring the role of CEACAM gene conversion among primates, we provide evidence that this process continues to shape CEACAM diversity within human populations. The three human CEACAM1 variants we test in our adhesin binding assay are part of a group of related CEACAM1 haplotypes that increase sequence similarity to CEACAM3 and/or CEACAM5 (Figure 6 – figure supplement 1). Extended haplotypes that increase similarly to CEACAM1 at both synonymous and nonsynonymous positions in the N-domain are also found for CEACAM3 and CEACAM5 in humans (Figure 6 – figure supplement 3 and 4). Indeed, haplotypes consisting of variants of putative recombination events are the most common non- reference alleles for CEACAM1, CEACAM3, and CEACAM5 (Figure 6A, Figure 6 – figure supplement 1-4). Variant sites in these proteins tend to lie along the protein binding interface of the N-domain and often impact residues known to influence adhesin recognition. The relationships between these different CEACAM haplotypes appears to be complex, as many different combinations of partial variant haplotypes exist for each CEACAM paralog. The haplotype structures we observe suggest these CEACAM variants are the result of one or more recombination events between paralogous sequences, likely followed by further recombination with the major CEACAM allele.

Important questions remain regarding the rapid evolution of a subset of primate CEACAM proteins. Among these questions is why CEACAM7 and CEACAM8 show similar patterns of evolution to bacterially antagonized CEACAMs despite no known instances of bacterial antagonism. The simplest explanation is that CEACAM7 and CEACAM8 are themselves the targets of as yet unidentified pathogen antagonists (Sintsova et al., 2015). Alternatively, their rapid evolution may reflect pressure to maintain binding with rapidly evolving CEACAMs (Gray-Owen and Blumberg, 2006; Skubitz and Skubitz, 2008), could merely be a result of their genomic proximity to rapidly evolving CEACAMs prone to gene conversion (Zid and Drouin, 2013) or could be the result of some as yet unknown evolutionary pressures.

Another intriguing aspect of rapid CEACAM evolution is the impact rapid divergence might have outside of interactions with pathogenic microbes. Given the extensive overlap of CEACAM binding sites among unrelated bacterial adhesins, the ramifications of rapid CEACAM evolution likely extend beyond the adhesins of pathogens to those of commensal and beneficial microbes as well. For commensal microbes which rely on these interaction surfaces, pathogen-driven evolution could significantly alter their ability to colonize the host. The impact of CEACAM divergence on composition of the host microbiome and/or the evolution of commensal strains warrants further investigation.

Studies of other “housekeeping” proteins targeted by pathogens have found that sites under positive selection typically do not overlap with sites involved in essential host functions (Barber and Elde, 2014; Demogines et al., 2013). This is clearly not the case for CEACAMs, where we observed extensive overlap between sites involved in host protein interactions, sites targeted by bacterial adhesins and sites undergoing rapid evolution (Figure 2B). How CEACAMs are able to rapidly evolve while maintaining their other essential host protein interactions remains a mystery. Future studies on CEACAM protein functions, interaction networks, and pathogen antagonism will likely clarify these outstanding questions regarding rapidly evolving CEACAMs.

Collectively our study provides evidence that repeated adaptation among primate CEACAMs has shaped host-specific cell adherence by diverse pathogenic bacteria. We find that over half of the CEACAM paralogs found in humans display signatures of positive selection across the primate lineage, localized primarily to the extracellular N-domain. We further discovered that rapid evolution of CEACAM N-domains has been facilitated by extensive “shuffling” of sequences between a subset of CEACAM paralogs through repeated gene conversion. The diversification of primate CEACAM N-domain sequences has likely had significant consequences for interactions between primates and bacteria. Consistent with observations across other primate species, we also provide evidence that gene conversion events impact bacterial pathogen recognition of CEACAMs in contemporary human populations. Together this work reveals how dynamic evolutionary processes have shaped bacterial-host associations with consequences for infectious disease susceptibility.

## MATERIALS AND METHODS

### Primate comparative genetics

#### Sequence identification

Orthologs for human CEACAM genes were identified through BLAST searches of primate reference genomes available through the NCBI BLAST webserver (Boratyn et al., 2013). Full length genomic regions for annotated human CEACAMs were used as query sequences. A full record of CEACAM orthologs identified and a partial record of BLAST results, including date accessed, query coverage and identity, as well as information on synteny, are listed in Supplementary files 4-6. Orthology was established by sequence identity, reciprocal best-BLAST hit, as well as intron structure and synteny. In total, we were able to extract 186 primate CEACAM sequences for analysis. We could not identify orthologs of every human CEACAM in every primate species, in some cases because of lineage specific gains and losses and in some cases likely because of incomplete genome assembly. As a result, the number of primate orthologs available for evolutionary analysis and phylogenetic reconstruction for each human CEACAM range from 11-19 (Supplementary file 4).

#### Sequence alignment & trimming

Orthologous protein coding sequences were extracted from CEACAM genes as follows. Multiple sequence alignments of the full-length gene were done using MAFFT alignment software as implemented in Geneious Prime 2020.2.2 with default settings. Alignments were manually corrected to correspond to human exon splice sites. Regions corresponding to human exons were then extracted, realigned, and minimally trimmed so all sequences were in-frame and orthologous codons aligned. So as not to exclude any protein coding regions from evolutionary analysis all human exons for a given CEACAM were concatenated and treated as a single protein coding sequence. Consequently, representations of CEACAM proteins in figures are not necessarily indicative of mature peptides, but rather represent all parts of the CEACAM protein that could potentially have been subject to positive selection. Gaps in alignments were removed for evolutionary analyses but were retained for tree building.

#### CEACAM3 exons

Almost all Old World monkey CEACAM3 genes analyzed had two extra exons annotated compared to humans. These exons are located between the exon encoding the N-domain and the transmembrane domain and are predicted by InterProScan (Quevillon et al., 2005), as implemented in Geneious Prime 2020.2.2, to encode the IgC-like domains typical of this region of CEACAM proteins. The majority of Old World monkeys have two exons annotated and all primates, including hominids, have strongly conserved sequences in this region, though hominids all encode premature stop codons. With the exception of the second IgC exon in colobus, these exons would allow for the translation of full length CEACAM proteins. While exon annotation differences between primate CEACAM genes is not unusual, the conservation of these sequences across primates containing a CEACAM3 gene, including in hominids where they are not annotated, was striking. To the best of our knowledge CEACAM3 transcripts for humans or other primates including either of these extra IgC domains have not been reported and indeed, the exon closest to the N-domain likely does not encode a functional protein in most hominids as a result of a premature stop codon. However, the strong conservation of these sequences across primates could indicate these exons encode functional protein segments in at least some species. For this reason, these exons and their orthologous sequences in hominids were included in downstream evolutionary analyses.

#### CEACAM5 trimming

The differences in number and likely arrangement of IgC domains in primate CEACAM5 orthologs prevented alignment of all full length CEACAM5 genes into a single multiple sequence alignment for extracting human orthologous protein coding sequences. Instead, sequences were first aligned in three groups; New World monkeys, leaf-eating monkeys (black-and-white colobus, black snub-nosed monkey and golden snub-nosed monkey), and the remaining Old World monkey sequences with the hominid sequences. There were enough similarities with human exons for orthologous exon sequences to be assigned and extracted for New World monkeys and the Old World monkey/Hominid group, but not for the leaf-eating monkeys group. For leaf-eating monkeys the predicted exons in common between species in this group were extracted. After extracting coding sequences for each group individually the extracted sequences were then aligned in a single multiple sequence alignment. However, the large gaps caused by missing IgC sequences relative to human CEACAM5 posed a problem for evolutionary analyses which require gaps to be removed from sequences prior to analysis. We were concerned that choices made regarding which sequences were removed would unduly influence the results of evolutionary analyses or result in lower coverage of the evolutionary history of the entire coding sequence. To account for this, three strategies of trimming alignment gaps were carried out and the results of each used in separate evolutionary analyses. For the first strategy every species whose sequence contained gaps corresponding to missing IgC domains was removed. These species were black-and-white colobus, black snub-nosed monkey, golden snub-nosed monkey, drill, sooty mangabey, and common marmoset. This resulted in the longest sequence for analysis (2 Kb) including 6 predicted IgC domains, but the smallest number of species represented (12). In the second strategy primate sequences with gaps corresponding to the largest number of missing IgC domains (four) were removed, while those with only two missing domains were retained, and the alignment region containing the sequence gap caused by the missing domains removed, giving a smaller alignment (1.4 Kb, with four IgC domains), but more species (16). For this strategy sooty mangabey, and common marmoset were removed from the analysis. For the third strategy all species for which complete CEACAM5 gene sequences could be identified were retained and all gaps corresponding to missing IgC domains removed. This gave the smallest sequence (0.9 Kb, retaining two IgC domains), but provided the largest number of represented species (18). Evolutionary analyses for these strategies are included in Figure 2 – figure supplement 1, Supplementary file 1 and Supplementary file 2.

#### Alignment comparison between MAFFT and MUSCLE

To confirm that our alignment method was not biasing the assignment of orthology of coding sequences to human exons, we compared the results of alignments of extracted exons using MAFFT (Katoh and Standley, 2013) and the alternative program MUSCLE (Edgar, 2004), both as implemented in Geneious Prime 2020.2.2. With the exceptions of CEACAM7 and CEACAM5 there were no drastic changes between alignments performed using MAFFT and those done using MUSCLE. Upon inspection the discrepancy between MAFFT and MUSCLE alignments for CEACAM7 could be attributed to an approximately 7 Kb insertion in the orangutan CEACAM7 gene relative to all other primates. Upon removing this insertion alignments with both MAFFT and MUSCLE were in agreement. Discrepancies between alignments of CEACAM5 with MAFFT and MUSCLE were due to differences in how the programs aligned sequences corresponding to IgC domains, likely as a result of differences in the number and possibly the arrangement of sequences coding for IgC domains between primates. MAFFT and MUSCLE alignments were carried out for each of the three different trimmed versions of CEACAM5 (see above) and each set of sequence alignments was tested using each of the evolutionary analysis methods. All other evolutionary analyses were carried out using sequences trimmed according to MAFFT alignments.

The results of CEACAM5 evolutionary analyses were largely similar regardless of which alignment or trimming method was employed, identifying similar patterns of selection (sites under selection concentrated in the N-domain) and many of the same sites under selection. Results presented in the paper are for dataset 1 (ds1) which contains the largest number of domains and using the MAFFT alignment to match the method used for other CEACAM analyses presented. Results for alternative CEACAM5 trimming and alignment methods are included in Figure 2 – figure supplement 1, Supplementary file 1 and Supplementary file 2.

#### Bonobo CEACAM1 N-domain sequence verification

Bonobo genomic DNA was not available for direct sequencing of CEACAM1, so currently available bonobo genome sequence data was used for sequence verification. While the genome assembly from which bonobo CEACAM sequences were identified for evolutionary analyses did not have reads available, a more recent assembly of a different bonobo individual became available during the course of this study which did deposit sequencing reads along with a *de novo* genome assembly (Mao et al., 2021). The CEACAM1 genomic region of the newer assembly was 99% identical to the older version while the coding sequences differ at only a single nucleotide outside of the N-domain. Furthermore, examining the reads used to assemble the newer genome we confirmed that multiple reads covered the length of the bonobo CEACAM1 N-domain and included the highly diverged nucleotides of the binding region in contiguous reads. Additionally, we examined CEACAM1 sequences for the thirteen bonobo individuals sequenced as part of the Great Apes Genome Project (Prado-Martinez et al., 2013). Genomes for these individuals were constructed using a reference based assembly method to the human genome. The assembled sequences largely supported the highly diverged N-domain seen in the reference genome; however there was a 31 bp region that was identical to the human CEACAM1 sequence rather than the two *de novo* bonobo sequences. Examining reads from these individuals failed to support human sequences at this position and in fact supported the more divergent sequence seen in the bonobo *de novo* assemblies. Nucleotide BLAST searches on the NCBI webserver for bonobo N-domain sequences were performed with query sequences searching against the RefSeq Genome Database (refseq_genomes) for the organism groups “Homo/Pan/Gorilla groups” (taxid:207598) and “Primates” (taxid: 9443), while excluding “bonobos” (taxid:9597).

#### Identification of human CEACAM N-domain variation

Human haplotype data for CEACAM1, CEACAM3, CEACAM5, CEACAM6, CEACAM7, and CEACAM8, available through the International Genome Sample Resource (https://www.internationalgenome.org/) was accessed through the Ensemble genome browser (https://www.ensembl.org/). For each CEACAM the haplotypes identified for the Matched Annotation from NCBI and EMBL-EBI (MANE) Select v0.92 transcript were used. All coding sequence haplotypes for the MANE Select transcript were downloaded and analyzed in excel as well as in R using custom scripts (Figure 6 – source data 1, Figure 6 – figure supplement 3 – source data 1, and Figure 6 – figure supplement 4 – source data 1).

### Phylogenetic analyses

#### PAML/FUBAR/MEME/GARD

Evolutionary analyses were performed individually for each group of human CEACAM coding sequence orthologs. Only CEACAM21 was excluded from evolutionary analyses, since it was found only in hominid genomes and has likely been lost in the pan lineage (Supplementary files 4 and 5) resulting in only three closely related sequences being available for comparison, insufficient for robust phylogenetic based evolutionary analysis. CEACAM21 sequences were included in subsequent phylogenetic reconstructions.

CEACAM coding sequences and specific amino acids were tested for evidence of positive selection using the PAML NS sites program under the codon model F3x4 (Yang, 2007). A relevant primate species tree, based on primate species relationships detailed in Pecon-Slattery 2014 (Pecon-Slattery, 2014), was provided for each analysis. To determine the likelihood a gene was evolving under positive selection, likelihood ratio tests were performed comparing the models of selection M1&M2 as well as M7&M8 (Supplementary file 1) (Bielawski, 2013). Sites evolving under positive selection were identified by PAML using the Bayes Empirical Bayes analysis as implemented in the NS sites package for evolutionary Model 2, which has been shown to be more robust to error due to recombination than the alternative, Model 8, when identifying sites under selection (Anisimova et al., 2003). In addition, sites under selection were identified (Supplementary file 2) using the HyPhy package programs FUBAR and MEME (Murrell et al., 2013, 2012) as implemented on the Datamonkey web servers (www.datamonkey.org and <u>classic.datamonkey.org</u> respectively) (Delport et al., 2010; Kosakovsky Pond and Frost, 2005; Pond et al., 2005; Weaver et al., 2018). For FUBAR and initial MEME analyses, species trees of the relevant primates were provided to inform analyses of evolution. HyPhy GARD analyses (<u>classic.datamonkey.org</u>) were used to identify evidence of recombination and the number and approximate locations of breakpoints (Kosakovsky Pond et al., 2006). When GARD detected evidence of recombination, updated GARD informed phylogenies were used for MEME analyses to account for errors in calling sites under selection due to recombination. Prior to running MEME and GARD analyses the “automatic model selection tool” provided by <u>classic.datamonkey.org</u> was used to determine the most appropriate model of selection under which to run analyses. For PAML, sites with posterior probability >0.95 were considered to have strong support to be evolving under positive selection (Yang et al., 2005), while >0.9 posterior probability supported sites found by FUBAR (Murrell et al., 2013) and p-values ≤0.05 supported sites found by MEME (Murrell et al., 2012).

#### Tree building

Phylogenetic trees were constructed using our panel of primate CEACAM coding sequences identified as described above. Multiple sequence alignments on which tree constructions were based were done by translation alignment using default settings of the MAFFT sequence alignment software as implemented in Geneious Prime 2020.2.2. For domain specific phylogenetic reconstruction domains were identified using InterProScan (Quevillon et al., 2005) in Geneious Prime. Assignments for immunoglobulin-like domains, that is the IgV-like (N-domain) and IgC domains were based on predictions by the Superfamily database (Wilson et al., 2009) and cytoplasmic domain assignments were based on the PHOBIUS database (Käll et al., 2004). Transmembrane domains were excluded from analyses due to their particularly small sequence length, which can make tree building unreliable due to limited phylogenetically informative sites. Indeed, relatively short sequence lengths for the other domains, typically around 300 bps or less, along with often high sequence similarity likely decreased the reliability and statistical support for our domain trees. However, even with these limitations in many cases relationships between domains were resolved with high bootstrap support, particularly for peripheral nodes and clades and for CEACAMs not found to be evolving rapidly. Phylogenetic reconstructions were done using the PhyML 3.0 web browser (http://www.atgc-montpellier.fr/phyml/) with default settings and confidence testing by 1000 bootstrap replicates (Guindon et al., 2010).

#### Data visualization

Visualization of evolutionary analyses, phylogenetic trees, sequence identity, and haplotype frequencies was done in R (R Core Team, 2019) using the R packages BiocManager (Morgan, 2019), treeio (Wang et al., 2019), ggplot2 (Wickham, 2016), ggtree (Yu et al., 2018, 2017), evobiR (Blackmon and Adams, 2015), and ggforce (Pedersen, 2021). Protein structures were visualized using the UCSF Chimera package version 1.13.1. Chimera is developed by the Resource for Biocomputing, Visualization, and Informatics at the University of California, San Francisco (supported by NIGMS P41-GM103311) (Pettersen et al., 2004).

### CEACAM1 Binding Assays

#### Recombinant CEACAM1 expression plasmid construction

Plasmids encoding primate CEACAM1 N-domains were constructed by assembly PCR and ligation independent cloning (LIC) into the pcDNA3 GFP LIC vector (6D) (a gift from Scott Gradia; Addgene plasmid #30127). A detailed description of the assembly PCRs is provided in the Supplementary file 7 and the DNA oligomers and templates are described in Supplementary files 8 and 9. Briefly, oligonucleotides were designed to assemble expression cassettes containing the human IgƘ signal sequence followed by a primate CEACAM1 N-terminal domain, and finally a STREPII tag and Tobacco Etch Virus (TEV) protease site. LIC cloning of the primate CEACAM1 N-terminal expression cassettes into pcDNA2 GFP LIC (6D) was performed following the protocol provided by the California Institute for Quantitative Biosciences at Berkeley (https://qb3.berkeley.edu/facility/qb3-macrolab/projects/lic-cloning-protocol/). Mutations to introduce bonobo CEACAM1 residues and population variants into the human CEACAM1 reference as well human CEACAM1 residues into the bonobo CEACAM1 reference sequence were done by site directed mutagenesis using mutation specific primers designed using the Agilent QuikChange Primer Design tool (https://www.chem.agilent.com/store/primerDesignProgram.jsp), then transformed into One Shot™ Top10 chemically competent cells for amplification and sequence verification. Plasmids were extracted for further use using the ZymoPURE™ II Plasmid Maxiprep kit.

Recombinant CEACAM1 expression plasmids were transfected into Human HEK293T cells using the Lipofectamine™ 3000 transfection kit following manufacturers instructions. Two days post transfection cell supernatant was collected and filter sterilized and cells were collected and lysed. Expression of proteins was confirmed by western blotting.

#### Bacterial strains & culture

*H. pylori* strains G27 (Baltrus et al., 2009) J99 (Alm et al., 1999), Tx30a (ATCC^®^ 51932), and the G27 HopQ deletion strain (*omp27::cat-sacB* in NSH57) (Yang et al., 2019) were cultured microaerobically at 37°C on Columbia agar plates supplemented with 5% horse blood 0.2% beta cyclodextrin, 0.01% amphotericin B, and 0.02% vancomycin. To assay binding between recombinant primate CEACAM1 N-domain proteins and *H. pylori* strains, *H. pylori* strains were grown for two to five days on solid media, collected and suspended in Brain Heart Infusion Media. 500uL of bacterial suspension were then incubated with 100uL of CEACAM protein for thirty minutes, rotating on a nutator. Bacteria were then washed twice with cold PBS. Samples to be visualized by western blotting were pelleted and resuspended in 1x Laemmli Buffer. Samples to be examined by flow cytometry were suspended in 0.5-1 mL of PBS.

The use of *Escherichia coli* to express MS11 and VP1 *Neisseria gonorrhoeae* Opa proteins was described previously (Roth et al., 2013). For this project plasmids expressing Opa proteins, Opa52 (Kupsch et al., 1993) and Opa74 (Roth et al., 2013), were synthesized in the pET-28a vector background by GeneScript. Synthesizing Opa expression plasmids bypassed the subcloning described in previous works that allowed outer membrane expression, so an N-terminal signal sequence from the OMP A protein, native to the outer membrane of *E. coli,* was added by the manufacturer to express Opa proteins on the outer membrane of *E. coli*. NcoI and HindIII restriction sites were used to add OMP A and Opa sequences to the pET-28a plasmid. Opa expression vectors were transformed into *E. coli* DH5α cells for maintenance, replication and sequence verification. Plasmids were extracted for further use using the Zymo Research Zyppy™ Plasmid miniprep kit. For pulldown experiments Opa expression plasmids were transformed into BL21(DE3) *E. coli* cells to allow for inducible expression of Opa proteins. Cells were grown to an optical density of OD600 0.4-0.6, then IPTG (Isopropyl β- d-1-thiogalactopyranoside) was added to a concentration of 100mM to induce expression of Opa proteins. Bacterial cells were left to induce for three hours at 37°C. For pulldown assays 300μL of induced *E. coli* cell culture was incubated with 100μL of CEACAM1 protein construct as processed as described for *H. pylori*. All *E. coli* cells were cultured at 37°C in LB (Luria-Bertani) broth.

#### Western blotting and flow cytometry

Pulldown experiments assayed by western blotting were visualized using a commercially available mixture of Mouse ⍺-GFP clones 7.1 and 13.1 (Sigma-Aldrich) at a dilution of 1:10^3^ for the primary antibody incubation followed by secondary incubation with goat ⍺-mouse conjugated to horseradish peroxidase (1:10^4^ dilution) (Jackson ImmunoResearch) and visualized by WesternBright™ ECL HRP Substrate (Thomas Scientific). For pulldowns visualized by western blotting CEACAM1 protein input samples were prepared by mixing 20uL of protein with 20uL 2x Laemmli then boiled at 95°C for five minutes and centrifuged at max speed for five minutes, before visualization by western blotting along side pulldown samples. GFP fluorescence of primate CEACAM1 constructs bound to *H. pylori* strain G27 was also measured by flow cytometry, with 10,000 events per sample measured. Flow cytometry data was analyzed using FlowJo v10.5.3. Western blots and flow cytometry experiments depicted are representative of at least three independent replicates performed at different times.

## AUTHOR CONTRIBUTIONS

E.P.B. and M.F.B. conceived the study with input from K.M.K. Phylogenetic analyses were performed by E.P.B. with assistance from K.M.K. CEACAM-adhesin binding experiments were performed by E.P.B. with assistance from R.S., E.B., C.K.G., and W.B. Original manuscript was prepared by E.P.B. with input from M.F.B. All authors contributed to editing of the original manuscript.

## ACKNOWLEDGMENTS

We are grateful to members of the Barber lab, especially Caitlin Kowalski, for helpful discussions and feedback on the manuscript. This work was supported by National Institutes of Health grants R35GM133652 (to M.F.B.) and F32AI147565 (to E.P.B). We thank Karen Guillemin for strains and assistance with *H. pylori* as well as Nina Salama for sharing *H. pylori* strains. We thank Andrew Kern and CJ Battey for helpful discussions and assistance with population genetic analyses. We thank Nels Elde for thoughtful discussions related to this work. We thank Jacob Laser for early assistance with CEACAM phylogenetics. We also thank Scott Gradia for the gift of vector pcDNA3 GFP LIC.

## COMPETING INTERESTS STATEMENT

The authors declare no competing interests.

## FIGURE SUPPLEMENTS

**Figure 2 – figure supplement 1.**
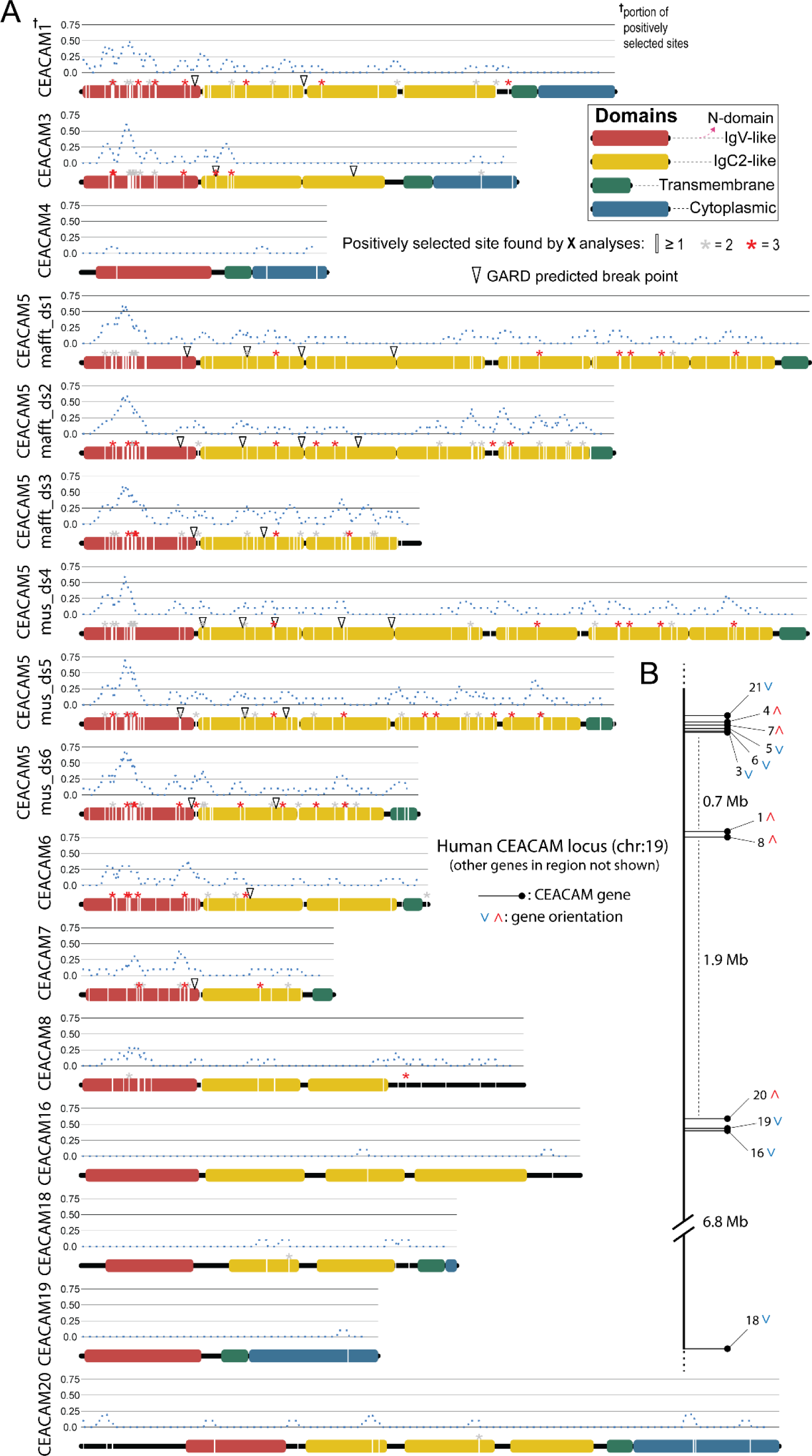
Primate CEACAM evolutionary analysis summary. Sites with elevated dN/dS in all human CEACAM proteins. A) Sites in CEACAM proteins identified as evolving rapidly in specific domains by one (white line), two (gray asterisks) or three (red asterisks) evolutionary analyses. Dotted blue line indicates the proportion of sites identified as evolving rapidly across a ten amino acid sliding window. Open triangles show GARD predictions of the approximate locations of recombination breakpoints. B) Location of human CEACAM genes along chromosome 19. Other genes on chromosome 19 are not shown.

**Figure 3 – figure supplement 1.**
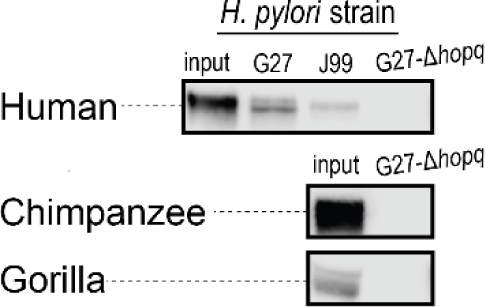
*H. pylori* G27 *Δhopq* pulldown Binding assay to assess interactions between *H. pylori* strain G27 *Δhopq* and GFP-tagged CEACAM1 N- domain constructs for human, chimpanzee, and gorilla, by pulldown experiments and visualization by western blot. Figure 3 – figure supplement 1 – source data 1. Raw and labeled western blot images for Figure 3 – figure supplement 1

**Figure 4 – figure supplement 1.**
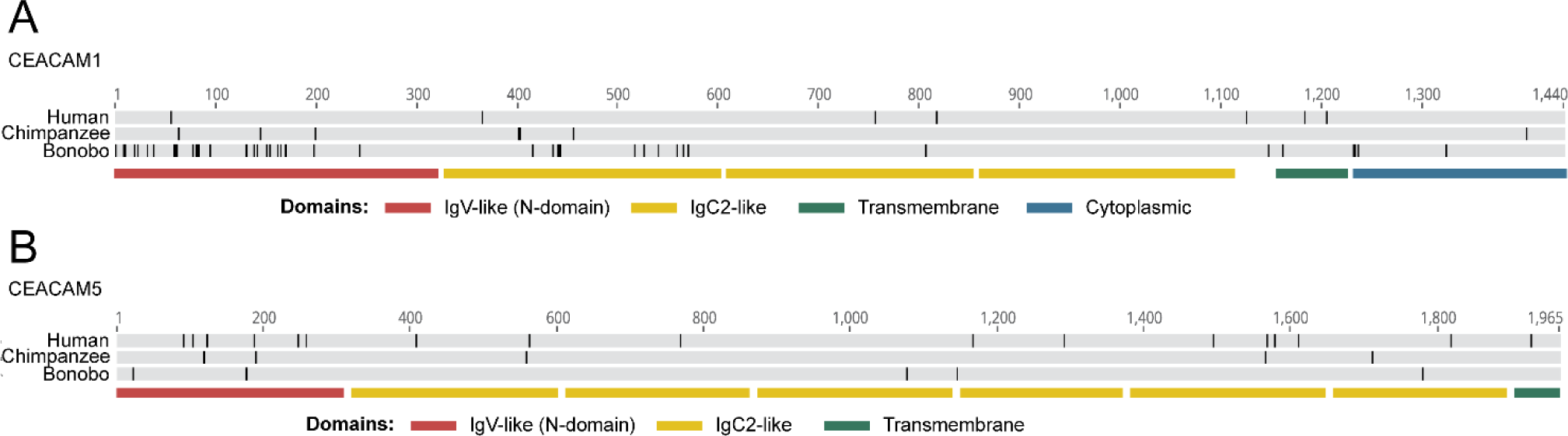
Alignment of Human-Pan CEACAM sequences. Bonobo CEACAM sequence alignments. Human, chimpanzee and bonobo CEACAM1 (A) and CEACAM5 (B) alignments by MAFFT translation alignment implemented in Geneious Prime 2020.2.2. Black lines mark differences from consensus. Lower lines show location of CEACAM domains.

**Figure 4 – figure supplement 2.**
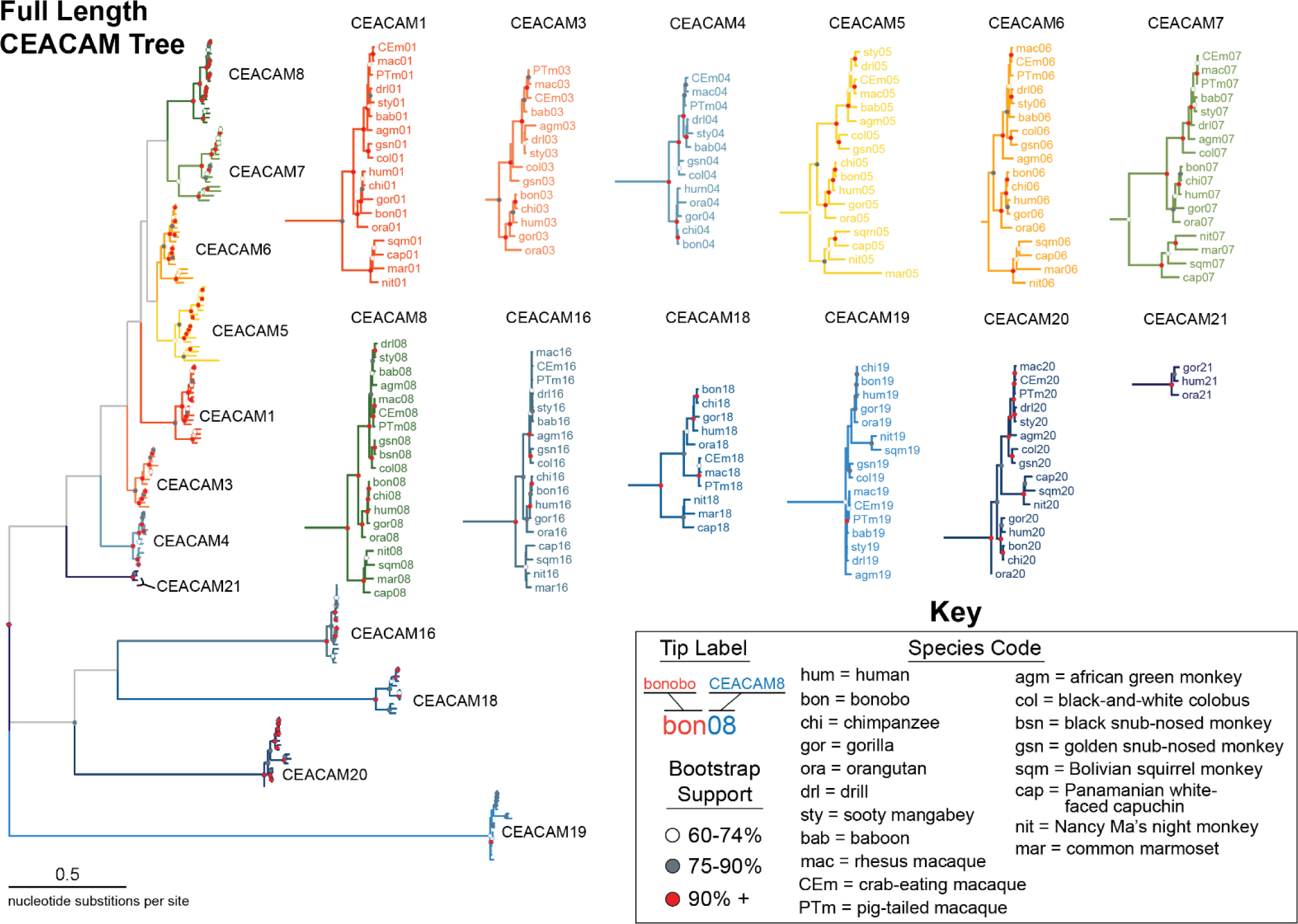
Expanded full length CEACAM tree. Maximum likelihood-based phylogeny of full length CEACAM protein coding sequences as represented in Figure 4A, but with clades expanded. Clades encompassing individual CEACAM orthologs are shown isolated and expanded.

**Figure 4 – figure supplement 3.**
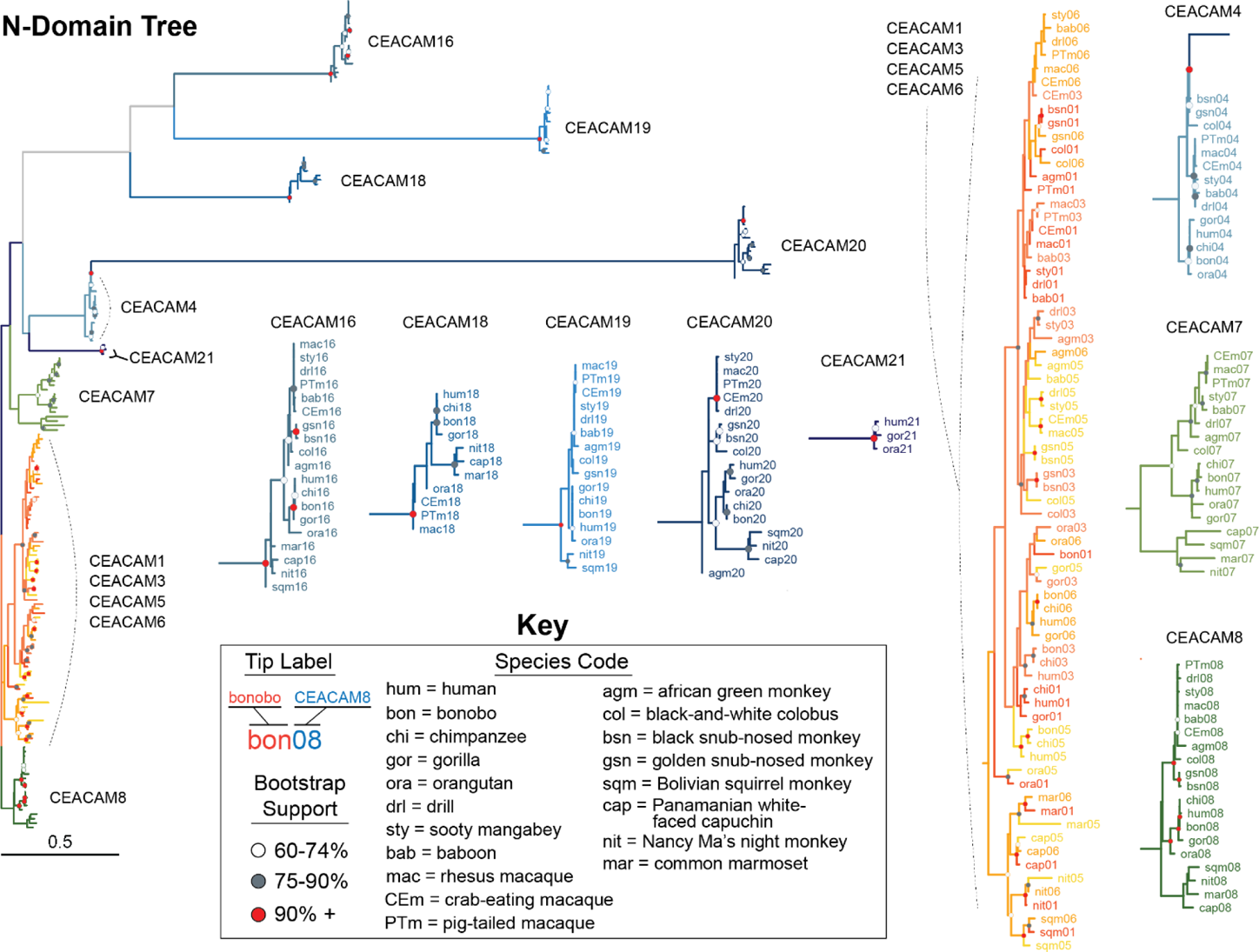
Expanded CEACAM N-domain tree. Maximum likelihood-based phylogeny of CEACAM IgV-like (N-domain) sequences as represented in Figure 4B, but with clades expanded. Clades encompassing individual CEACAM orthologs along with the CEACAM1, CEACAM3, CEACAM5 and CEACAM6 clade are shown isolated and expanded.

**Figure 4 – figure supplement 4.**
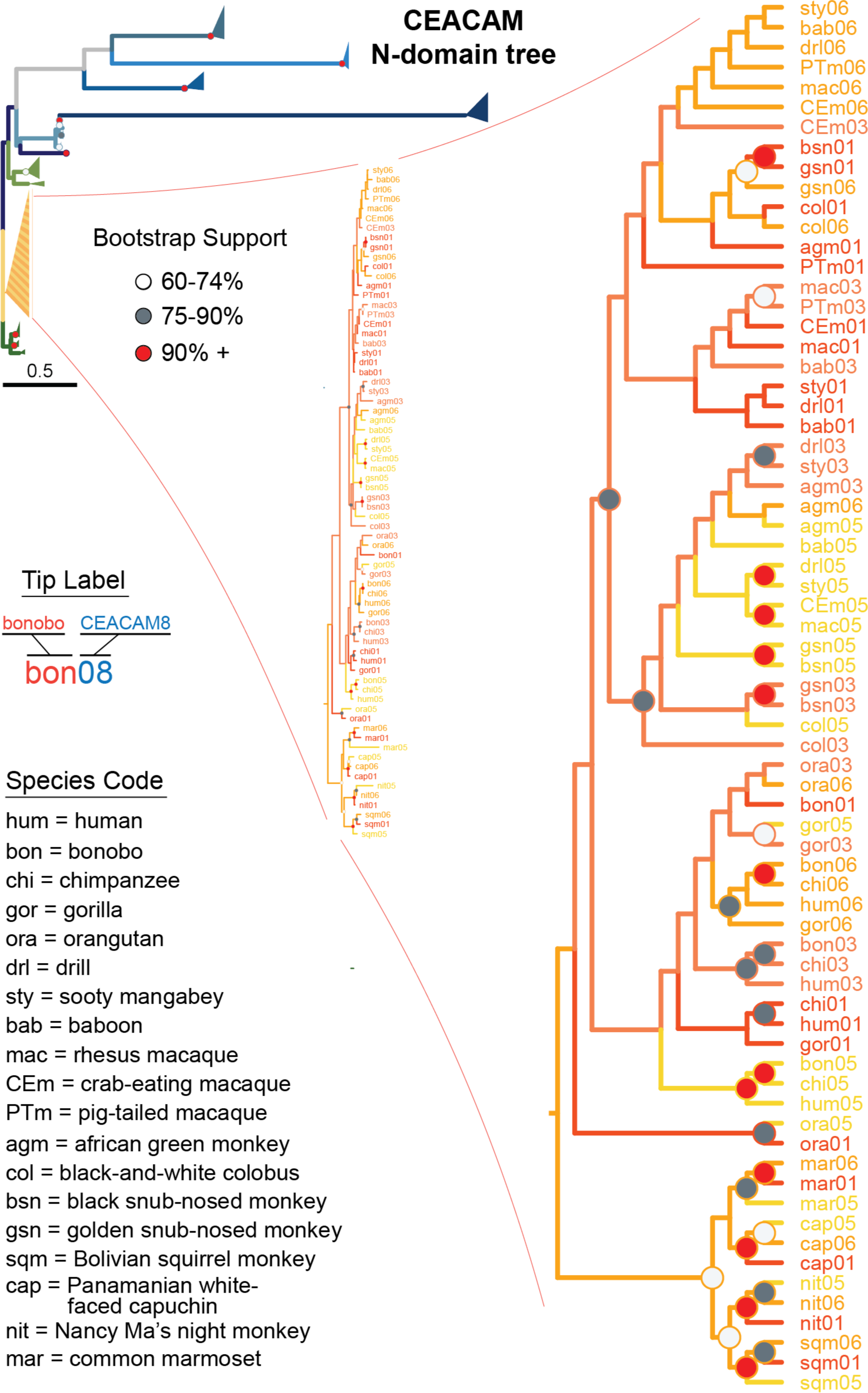
Expanded view of CEACAM1,3,5,6 N-domain clade. Expanded view of CEACAM1, CEACAM3, CEACAM5 and CEACAM6 clade from Figure 4B.

**Figure 4 – figure supplement 5.**
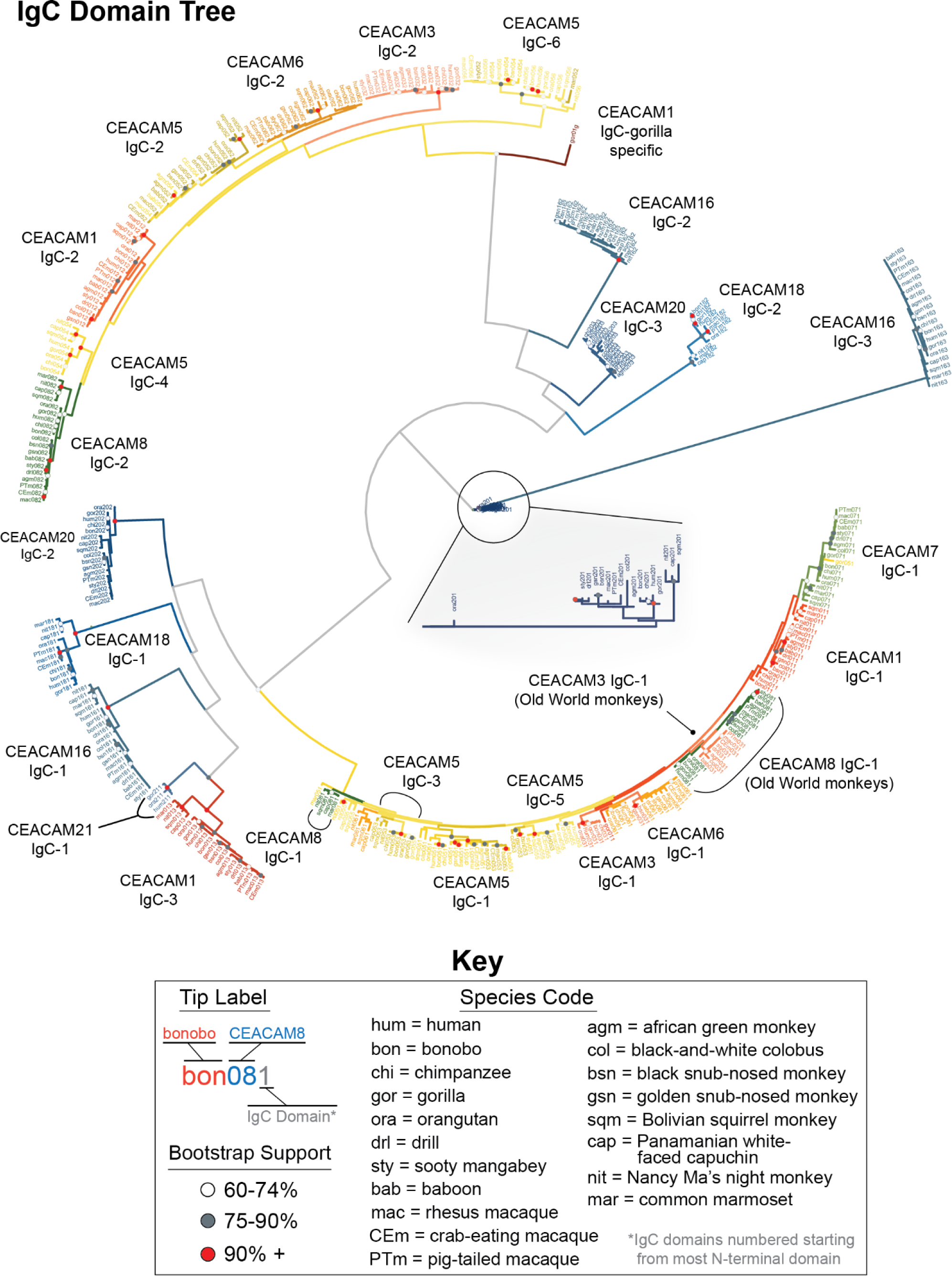
Expanded CEACAM IgC domains tree. Maximum likelihood-based phylogeny of CEACAM IgC-like domain sequences. Expanded view of CEACAM20 clade shown.

**Figure 4 – figure supplement 6.**
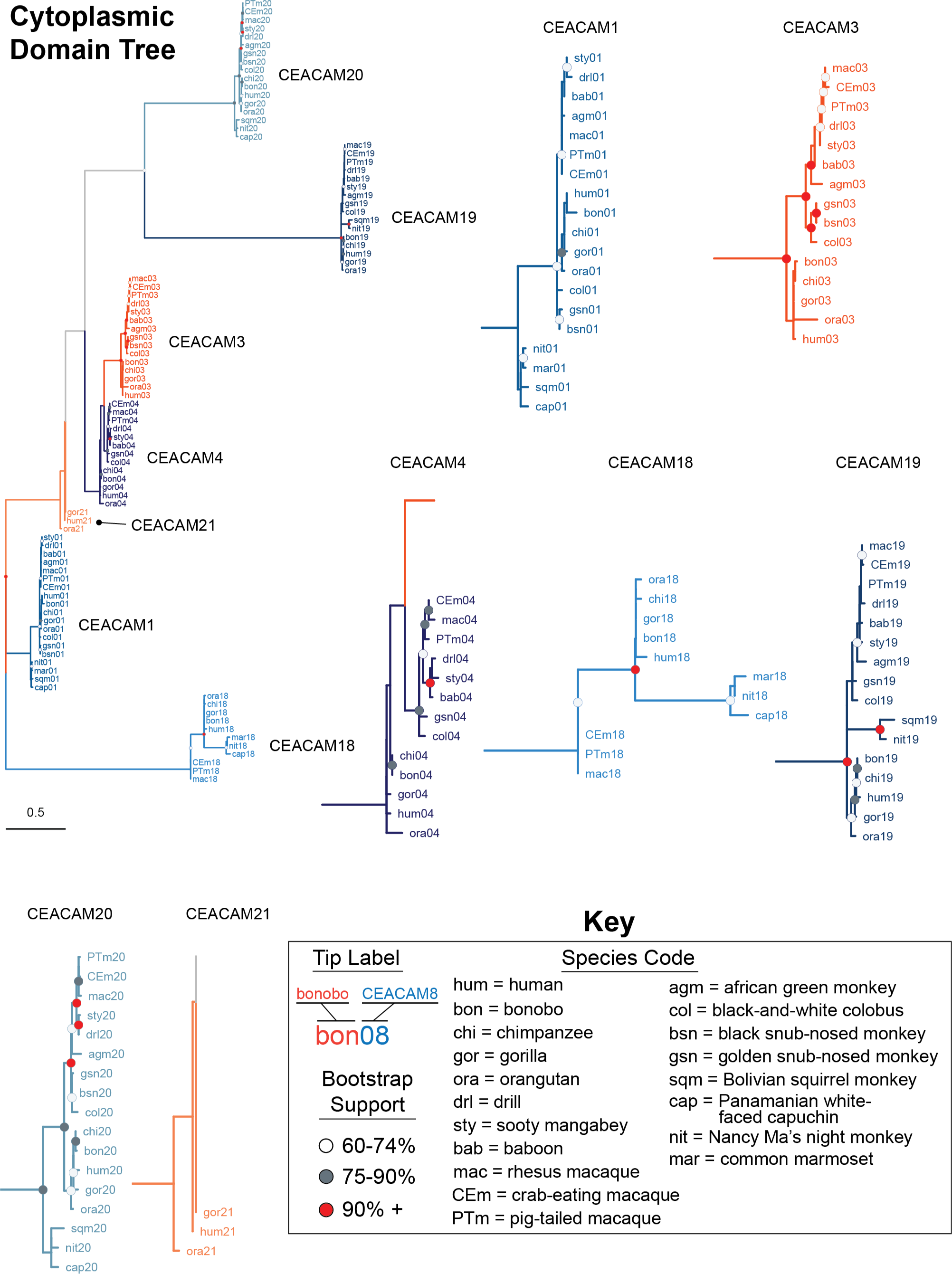
Expanded CEACAM cytoplasmic domain tree. Maximum likelihood-based phylogeny of CEACAM cytoplasmic domain sequences. Clades encompassing individual CEACAM orthologs are shown isolated and expanded.

**Figure 5 – figure supplement 1.**
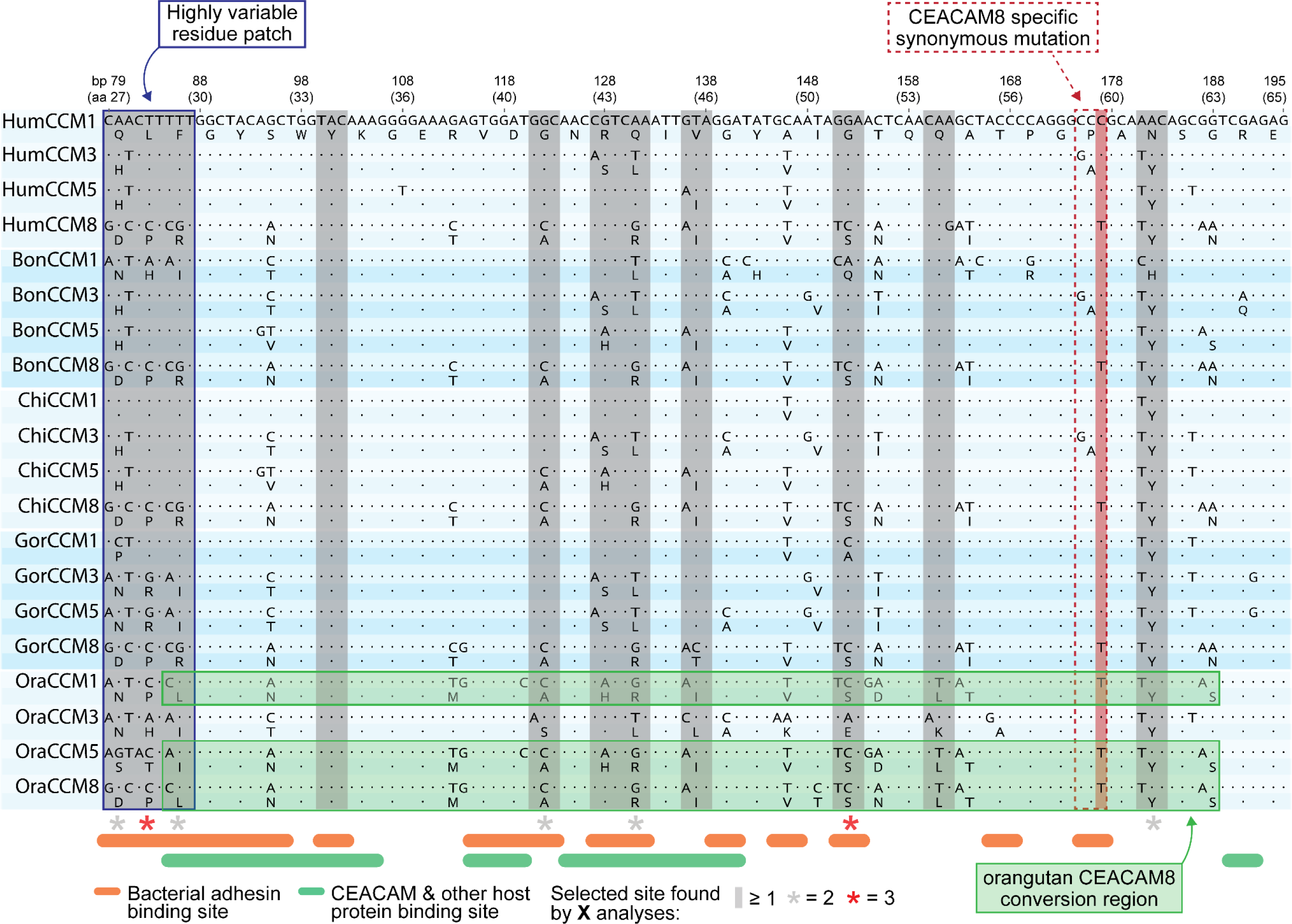
Alignment of rapidly evolving N-domain region in hominids. Multiple sequence alignment of CEACAM1, CEACAM3, CEACAM5 and CEACAM8 orthologs for human, bonobo, chimpanzee, gorilla, and orangutan. Translation of each nucleotide sequence is positioned on the line below. Sites known to influence adhesin and host protein binding (Supplementary file 3) are indicated as are sites identified as evolving under positive selection.

**Figure 6 – figure supplement 1.**
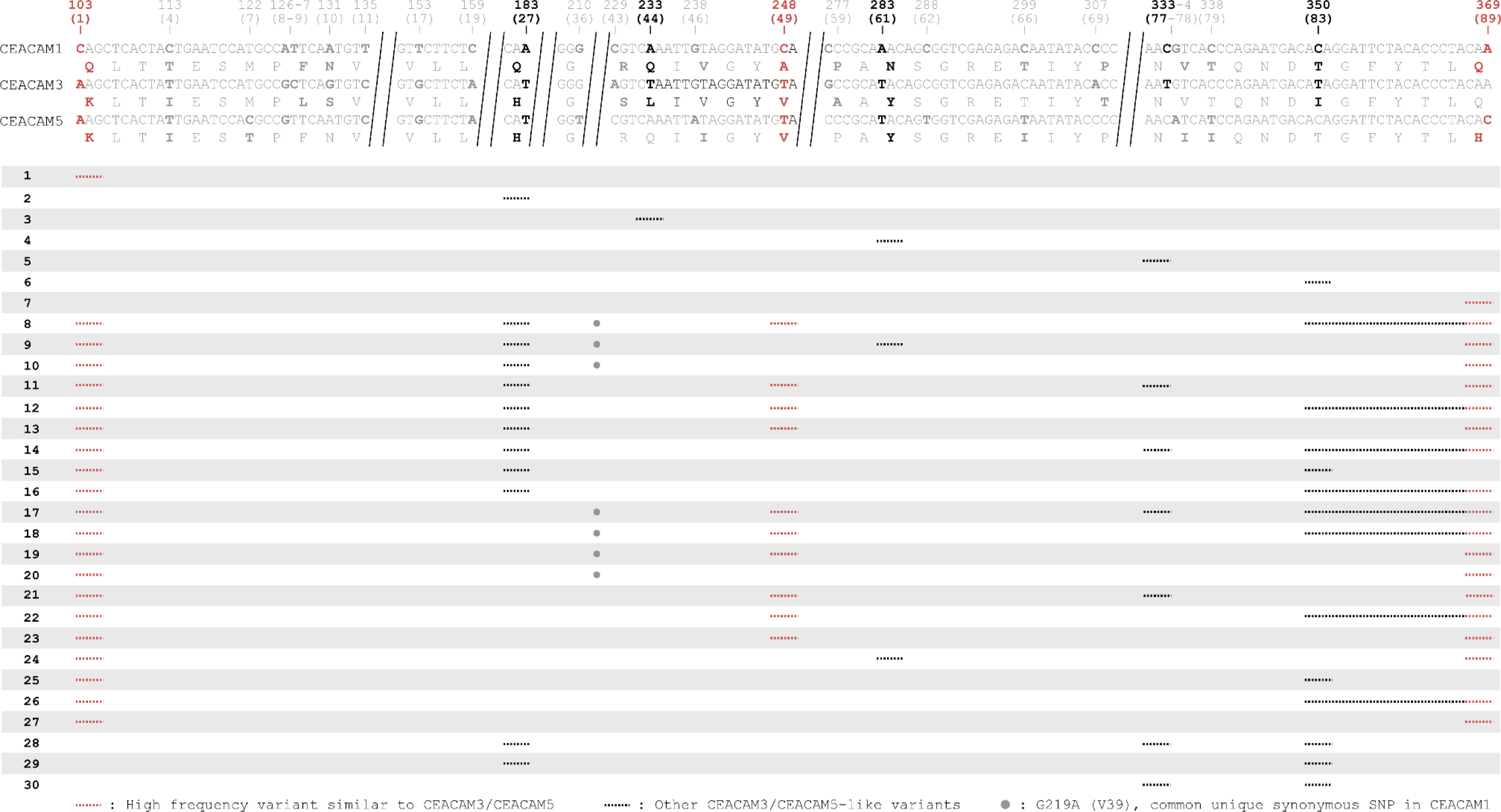
Human CEACAM-like CEACAM1 haplotypes. Other CEACAM-like human CEACAM1 haplotypes. Alignment of human CEACAM1, CEACAM3 and CEACAM5 N-domain reference nucleotide sequences with amino acid translations below. Long invariable alignment regions were removed. Sites that differ in CEACAM3 or CEACAM5 relative to CEACAM1 are bolded. Sites found in variant CEACAM1 haplotypes are in black. Changes that encode the high frequency variants Q1K, A49V, and Q89H are in red. Below alignment each row is a unique human CEACAM1 N- domain haplotype. Lines indicate variant regions in CEACAM1. Only haplotypes that increase similarity to CEACAM3 or CEACAM5 are shown.

**Figure 6 – figure supplement 2.**
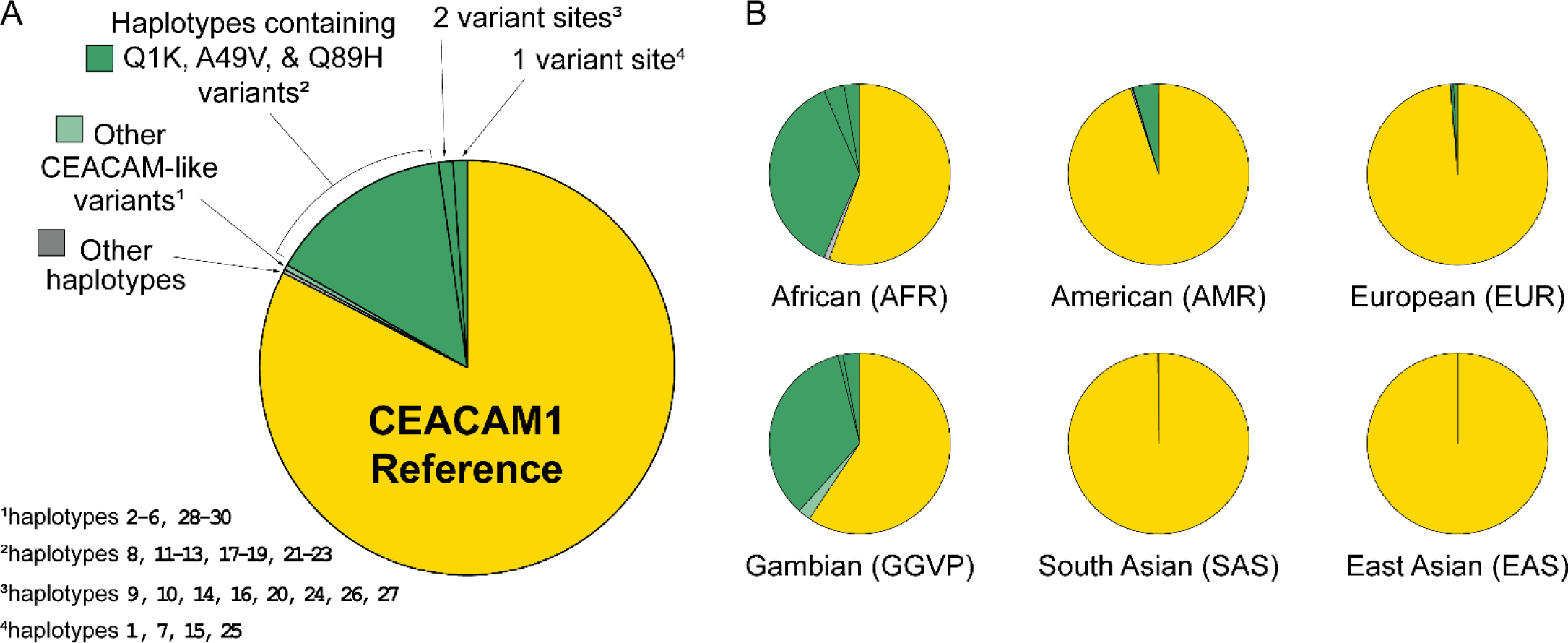
Human CEACAM1 haplotype frequencies. Frequency of variant human CEACAM1 haplotypes. A) Overall frequency of CEACAM1 variants Q1K, 449V, Q89H and other variant haplotypes in humans. The indicated CEACAM-like haplotypes are enumerated in Figure 6 – figure supplement 1. B) Frequency of CEACAM1 variants across different human populations.

**Figure 6 – figure supplement 3.**
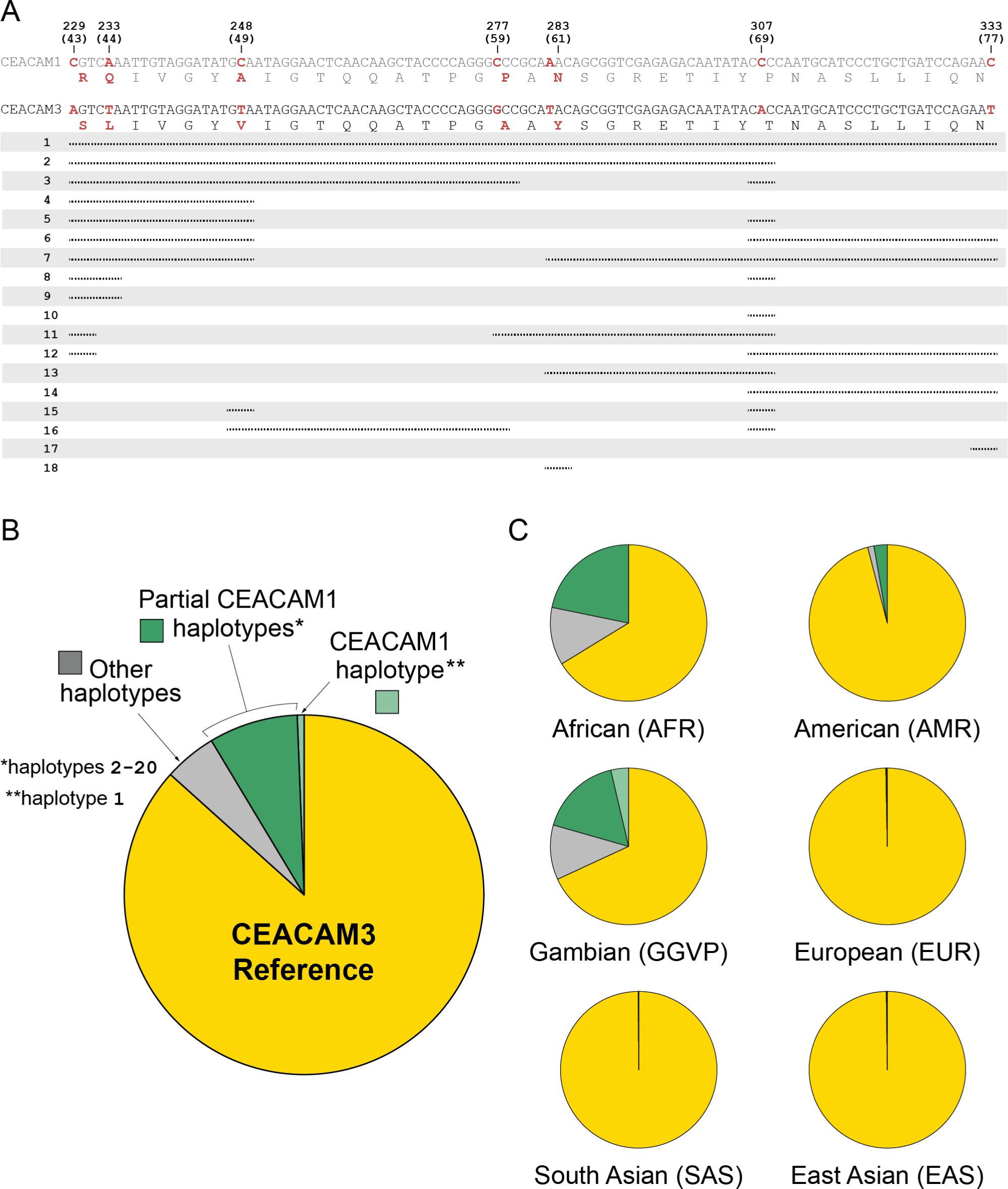
Human CEACAM3 variation. Human CEACAM1-like CEACAM3 haplotypes. A) Alignment of human CEACAM1 and CEACAM3 reference sequences. Disagreements are bolded in red with the amino acid translation below each sequence. Below alignment each row represents a unique human CEACAM3 haplotype. Lines indicate variant regions that match the human CEACAM1 reference sequence. Only haplotypes that increase similarity to the human CEACAM1 reference sequence are shown. B) Overall frequency of variant CEACAM3 haplotypes in humans. The CEACAM1-like haplotypes indicated are enumerated in panel A. C) Frequency of CEACAM3 variants across different human populations.

**Figure 6 – figure supplement 4.**
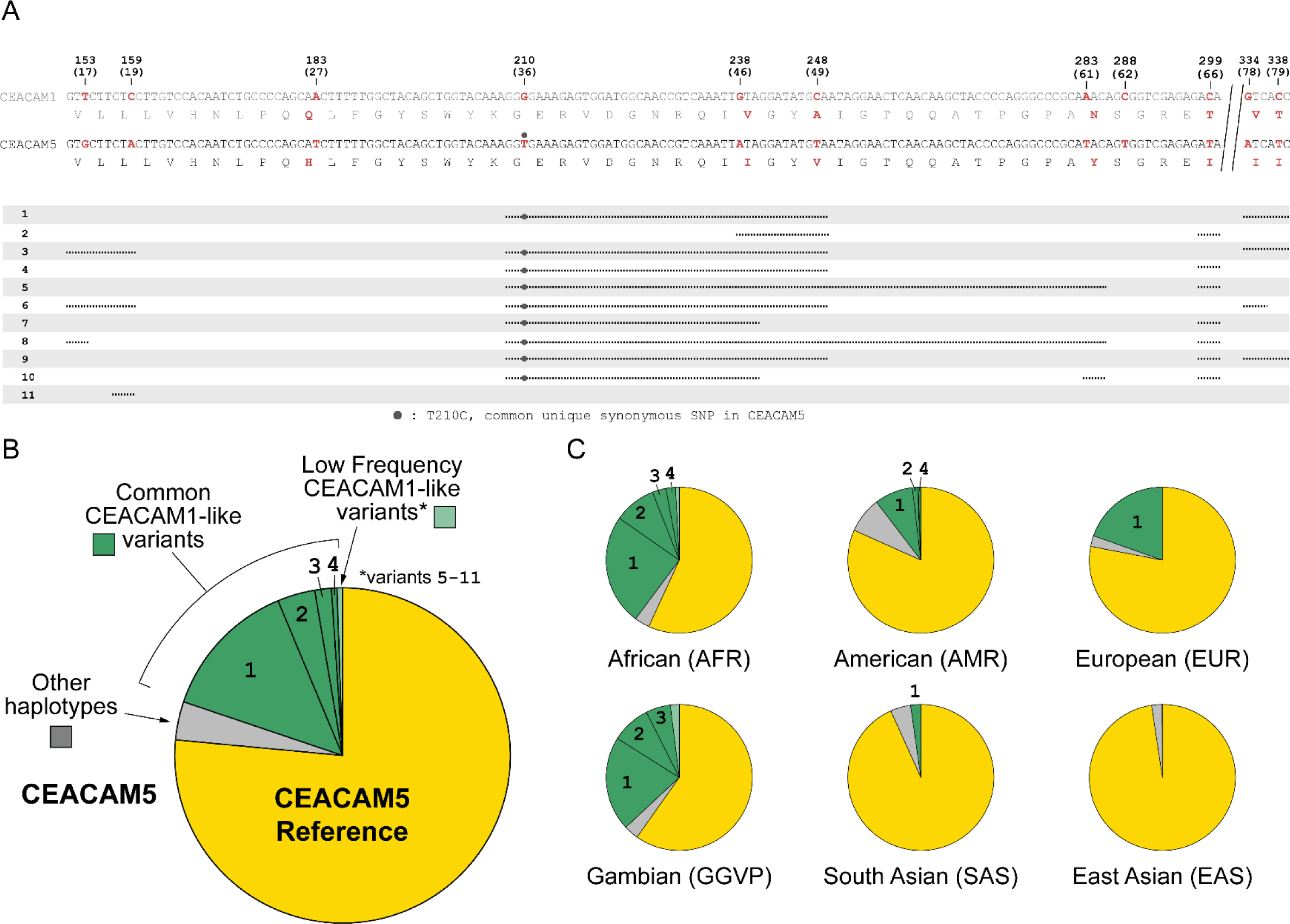
Human CEACAM5 variation. Human CEACAM1-like CEACAM5 haplotypes. A) Alignment of human CEACAM1 and CEACAM5 reference sequences. Disagreements are bolded in red with the amino acid translation below each sequence. Below alignment each row represents a unique human CEACAM5 haplotype. Lines indicate variant regions that match the human CEACAM1 reference sequence. Only haplotypes that increase similarity to the human CEACAM1 reference sequence are shown. B) Overall frequency of variant CEACAM5 haplotypes in humans. The CEACAM1-like haplotypes indicated are enumerated in panel A. C) Frequency of CEACAM5 variants across different human populations.

## SOURCE DATA

**Figure 3 – source data 1.** Raw and labeled western blot images for Figure 3A and flow cytometry data for Figure 3B.

**Figure 3 – figure supplement 1 – source data 1**. Raw and labeled western blot images for Figure 3 – figure supplement 1.

**Figure 6 – source data 1.** Data files and code for analyzing CEACAM1 haplotypes for Figure 6A and Figure 6 – figure supplement 2.

**Figure 6 – source data 2.** Raw and labeled western blot images for Figure 6C.

**Figure 6 – figure supplement 3 – source data 1.** Data files and code for analyzing CEACAM3 haplotypes for Figure 6 – figure supplement 3B and C.

**Figure 6 – figure supplement 4 – source data 1.** Data files and code for analyzing CEACAM5 haplotypes for Figure 6 – figure supplement 4B and C.

**Figure 5 – source data 1.** Raw and labeled western blot images for Figure 5C.

## SUPPLEMENTARY FILES

Supplementary file 1

Supplementary table 1: PAML NS sites tests of selection in primate CEACAMs

Supplementary file 2

Supplementary table 2: Sites identified as evolving under positive selection by evolutionary analyses and GARD predicted recombination breakpoints.

Supplementary file 3

Supplementary table 3: References for sites identified as contributing to CEACAM1 binding with host proteins and bacterial adhesins as well as the specific sites identified.

Supplementary file 4

Supplementary table 4: Table summarizing primate CEACAM sequences extracted for evolutionary analyses and phylogenetic reconstructions.

Supplementary file 5

Supplementary table 5: Table summarizing BLAST results, genome annotation, and sequence analyses used to identify human CEACAM orthologs in primates.

Supplementary file 6

Supplementary table 6: Additional notes on CEACAM sequences included and excluded from analyses

Supplementary file 7

Supplementary Methods

Supplementary file 8

Supplementary table 7: Table of oligomers, DNA templates and their order in assembly reactions used to assemble CEACAM1 N-domain expression plasmids.

Supplementary file 9

Supplementary table 8: Sources of template sequences for CEACAM1 and other plasmid components used for expression plasmid construction.

## Notes

### Competing Interest Statement

The authors have declared no competing interest.

### Summary of Updates

Additional control data was added for figures 5 and 6 and details for some methods were clarified.

